# Size-by-environment interactions: a neglected dimension of species’ responses to environmental variation

**DOI:** 10.1101/329771

**Authors:** Andrew T. Tredennick, Brittany J. Teller, Peter B. Adler, Giles Hooker, Stephen P. Ellner

**Affiliations:** Department of Wildland Resources and the Ecology Center, Utah State University, Logan, Utah; Department of Biology, Penn State University, University Park, Pennsylvania; Department of Biological Statistics and Computational Biology, Cornell University, Ithaca, New York; Department of Ecology and Evolutionary Biology, Cornell University, Ithaca, New York

**Keywords:** demography, demographic buffering, environmental variability, individual heterogeneity, individual size, integral projection model, population model

## Abstract

In both plant and animal systems, size can determine whether an individual survives and grows under different environmental conditions. However, it is less clear whether and when size-dependent responses to the environment affect population dynamics. Size-by-environment interactions create pathways for environmental fluctuations to influence population dynamics by allowing for negative covariation between sizes within vital rates (e.g., small and large individuals have negatively covarying survival rates) and/or size-dependent variability in a vital rate (e.g., survival of large individuals varies less than small individuals through time). Whether these phenomena affect population dynamics depends on how they are mediated by elasticities (they must affect the sizes and vital rates that matter) and their projected impacts will depend on model functional form (the impact of reduced variance depends on the relationship between the environment and vital rate). We demonstrate these ideas with an analysis of fifteen species from five semiarid plant communities. We find that size-by-environment interactions are common but do not impact long-term population dynamics. Size-by-environment interactions may yet be important for other species. Our approach can be applied to species in other ecosystems to determine if and how size-by-environment interactions allow them to cope with, or exploit, fluctuating environments.

## Introduction

The mechanisms by which environmental variability affects population dynamics are the focus of many longstanding questions in ecology: Does environmental variation have a negative effect on population persistence via stochastic extinctions (Stacey & Taper, 1992), or a positive effect via temporal niche partitioning (Chesson, 2000; Adler & Drake, 2008)? Do negative covariances among vital rates buffer populations against environmental variability (Jongejans *et al.*, 2010)? Can we use life history information to accurately predict population dynamics (Crone *et al.*, 2013)? Increases in climate variability and the need to forecast population trajectories in changing environments make these questions even more important. In size-structured populations, the answers to these questions could depend on how individual size affects responses to environmental variation.

There are two ways that size might influence a species’ environmental response. First, differently sized individuals can respond in different directions to environmental fluctuations. This would mean a “good” year for small individuals’ survival, for example, is a “bad” year for large individuals’ survival, and *vice versa*. We found no evidence of this in the literature, maybe because it has not been investigated by ecologists, or maybe it is unlikely that a bad year for one size class of a species can truly be a good year for another. Second, different size individuals can have different magnitudes of response to environmental conditions. In controlled experiments with weedy nodding thistle (*Carduus nutans*), warmer temperatures increased survival evenly across size classes, but smaller individuals disproportionately increased seed production in response to warming (Zhang *et al.*, 2011). Similarly, whether large or small yellow-bellied marmots (*Marmota flaviventris*) have a better chance at reproduction depends on environmental conditions (Ozgul *et al.*, 2010).

The environment is not just abiotic conditions, as biotic interactions also contribute to the total environment experienced by individuals. For example, plant defenses from herbivores typically change or develop over a plant’s lifespan, hence different life stages have different responses to time-varying herbivore pressure (Barton & Boege, 2017). Insofar as individual size and life stage are correlated, this implies different sized individuals will have different sensitivity to herbivory. Similarly, size-selective predation creates asymmetries in mortality rates between small and large individuals that can influence animal population dynamics (*e.g.,* Sprules, 1972; Hülsmann *et al.*, 2011).

These case studies are important for understanding the mechanisms by which individuals are affected by environmental conditions, but what remains unknown is whether and how size-by-environment interactions impact population dynamics. Understanding the effects of environmental conditions on individuals of different size could be essential for models with population structure, and ignoring differences due to size could bias predictions by over-emphasizing the response of the most numerous size classes. For example, the size distribution in many plant populations is skewed toward small individuals, but large individuals have the most impact on population dynamics (e.g., Dalgleish *et al.*, 2011). If survival rates of smaller individuals vary more year-to-year in response to environmental fluctuations than do large individuals, and small individuals comprise a large proportion of the dataset, statistical models with size-independent survival variation would be biased toward fitting the signal from small individuals. In this case, the interannual variance of survival for large individuals would be overestimated, which could strongly impact projected population dynamics. When different size classes within a population vary in sensitivity to environmental fluctuations, or respond to environmental conditions in opposite directions, models assuming size-independent environmental sensitivity will average those distinct responses and incorrectly predict temporal variation (or predict it correctly for the wrong reasons).

We propose that size-by-environment interactions may create additional pathways for environmental variation to influence population dynamics. They may allow for negative covariation among sizes for one vital rate (e.g., small and large individuals have negatively covarying survival rates) and/or reduced variance of a vital rate in one size class relative to another (e.g., survival of large individuals varies less than small individuals through time). Whether these phenomena affect population dynamics depends on how they are mediated by elasticities and how environmental variability is translated through individual-level responses up to population dynamics. Elasticities determine whether size-by-interactions matter because they must affect sizes and vital rates that have large contributions to population growth. How environmental variability impacts population dynamics through size-by-environment interactions depends on norms of reaction, which determine if reduced variance produces negative or positive effects. In this paper, we elaborate on these ideas and demonstrate them with an empirical analysis of fifteen species from five semiarid plant communities using a general analytical framework that can be easily applied to other time series of species’ abundances.

We begin by reviewing core concepts that link life history, environmental fluctuations, and population dynamics, and we introduce size as a new dimension to which these concepts apply. We then outline our statistical approach, including empirical examples from 15 perennial plant populations. Using the fitted statistical models, we probe model results and simulations of population dynamics to understand the impacts of size-by-environment interactions.

Our focus is on size-structured population models where size is a continous variable. Ageand stage-structured models often include (st)age-by-environment interactions implicitly by fitting time varying parameters for different (st)ages. For example, Hunter *et al.* (2010) parameterized a six-stage population model for polar bears (*Ursus maritimus*) where the extent of sea ice individually affected each stage transition. Thus, each stage is allowed to experience the environment in unique ways; the other extreme is represented by models which assume that environmental fluctuations affect all (st)ages the same way. What we propose below is a compromise that draws on the flexibility of generalized linear mixed effects models and their direct incorporation into integral projection models. Synthesis studies using stochastic matrix models with time-varying projection matrices have found that reduced variance of older individual’s survival can buffer populations against environmental variability (Morris *et al.*, 2008; Gamelon *et al.*, 2016). These findings set the stage for asking whether size-by-environment interactions provide similar opportunities for species to buffer themselves against environmental variability. Likewise, integral projection models with continuous size structure allow a greater opportunity to identify size-by-environment interactions because population structure is more finely resolved.

## Size-by-Environment Interactions and Population Dynamics

Size is one of the most important traits distinguishing individuals within a species, and demographic processes such as survival and growth are often largely determined by individual size (De Roos *et al.*, 2003). The relationships between size and vital rates are often assumed to be static through time. However, if vital rates do actually vary through time, this variability will affect population growth rates, often negatively (Lewontin & Cohen, 1969; Lande *et al.*, 2003). The negative effects of temporally variable vital rates suggests that species should evolve strategies to buffer themselves against environmental variability (Gillespie, 1977). This logic underlies the demographic buffering hypothesis, which states that species should evolve to have negatively covarying vital rates through time (Knops *et al.*, 2007) and/or have reduced variance of vital rates most important for fitness (Pfister, 1998).

Theoretical and empirical studies on demographic buffering have focused extensively on how vital rates vary through time without considering whether individual size mediates the temporal responses of species to environmental fluctuations. Previous studies have shown that different (st)ages of a population have unique responses to the environment (Coulson *et al.*, 2001) and that trade-offs among (st)age classes can lead to demographic buffering (Morris *et al.*, 2008; Gamelon *et al.*, 2016), suggesting that size-by-environment interactions may also be important. Conclusions from (st)age-structured models might also apply to size-structured populations, if size and (st)age are positively correlated.

Size-by-environment interactions provide a distinctive way for species to be buffered against environmental variability. In the presence of size-by-environment interactions, demographic buffering can arise through two similar mechanisms, as described above (Fig. 1). If different size classes within a population respond to the environment in opposite directions, such negative covariance could buffer population growth (path **a** in Fig. 1). This is similar to the idea of negative covariation between vital rates (e.g., survival and fecundity), but here the negative covariation is between size classes, within one vital rate. Buffering can also occur if the size classes that contribute most to population growth are the least sensitive to environmental conditions (path **b** in Fig. 1). This is similar to the idea of reduced variance in particular vital rates and/or (st)ages (Morris *et al.*, 2008; Gamelon *et al.*, 2016). Thus, individual size provides another dimension across which demographic buffering can occur.

**Figure 1:**
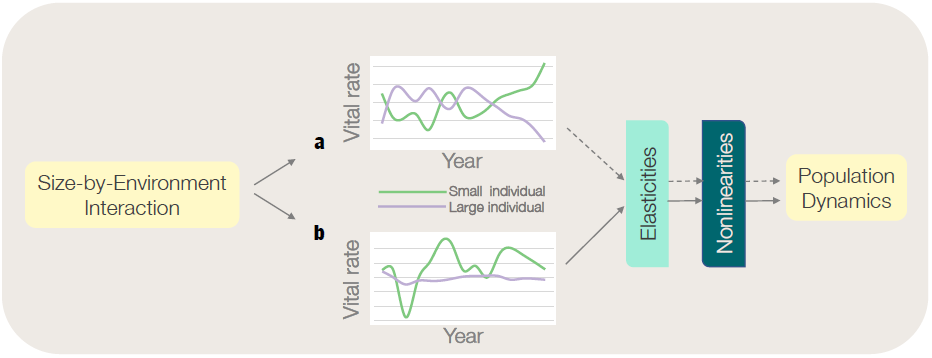
Conceptual overview of how size-by-environment interactions can impact population dynamics. If size-by-environment interactions are statistically important in vital rate regressions, there are two pathways by which they can impact population dynamics. In **a**, differently sized individuals have negatively covarying vital rates. For example, survival of small and large plants may be highest in different types of years. In **b**, the vital rate of one size varies less relative another sized individual. For example, small plant survival may vary more over time than large plant survival. Whether pathways **a** and **b** impact population dynamics depends on how they pass through the filters of elasticities and model functional form. The vital rates that negatively covary by size (**a**) must be the vital rates that are most elastic. Likewise, the sizes that vary more or less relative to other sizes must do so for the size-vital rate combinations that are most elastic. Last, whether pathway **b** has a positive or negative effect on population dynamics depends on the functional form of the vital rate’s relationship with size. For example, a concave-up relationship between size and growth rate would mean that large plants varying more than small plants is beneficial for population growth. Source code for figure insets: SizeByYear schematic.R.

However, size-by-environment interactions only matter for population dynamics if they affect sizes and vital rates that are important for population growth in stochastic environments. This rather obvious statement is implicit in empirical tests of the demographic buffering hypothesis. Typical tests involve calculating correlations between elasticities of vital rates and their temporal (co)variances (Morris & Doak, 2004). Considering size-by-environment interactions adds another layer: size and vital rate combinations must have high elasticity to impact population growth and fitness (Fig. 1).

The demographic buffering hypothesis assumes that environmental variation negatively affects population growth. If environmental variation was ‘good’, why would species evolve strategies to buffer themselves against it? This assumption is grounded in mathematical theory showing that short term variation in fitness reduces long term fitness when reaction norms are linear (Lande *et al.*, 2003; Morris *et al.*, 2008). If reaction norms are nonlinear, however, environmental variability can actually increase long term fitness (Drake, 2005; Koons *et al.*, 2009; Lawson *et al.*, 2015), a consequence of Jensen’s inequality (Jensen, 1906). In particular, concave-up and sigmoidal reaction norms provide the necessary conditions for environmental fluctuations to increase long term fitness and population growth. Environmental fluctuations can increase both the mean and the variance of annual population growth rate, consequently environmental fluctuations only benefit the long-run population growth rate λ_*S*_ (the geometric mean of a sequence a λ) if the increase in the mean overwhelms the negative effect of increasing the variance. Therefore, whether sizeby-environment interactions, which introduce temporal variability into the relationship between an individual’s size at time *t* and its size or probability of remaining alive at time *t* + 1, have positive or negative affects on population growth rate depends on the underlying reaction norm between individual-level vital rates and the environment. This reaction norm can be modeled as function of a measured environmental variable, or it can be estimated from random fluctuations in the response variable (e.g., individual survival). We take the latter approach in this paper, as described below.

## Detecting Size-by-Environment Interactions

### A flexible statistical approach

Detecting and quantifying size-by-environment interactions requires statistical estimates of interactions between size and time-varying environmental covariates. Variables related to weather conditions are obvious candidates for size-byclimate interactions, but it is difficult to quantify the effects of weather on demography or fitness. Identifying exactly which weather covariates, over which time scales, actually impact demographic rates is a thorny model selection problem. It requires longer time series than typically available to ecologists (Teller *et al.*, 2016; van de Pol *et al.*, 2016), or else it requires *a priori* assumptions about how weather should be aggregate into a few covariates, such as “spring rainfall” or “degree days.” Likewise, readily available weather data rarely exist at the fine spatial scales relevant to individual plants, meaning that measured weather covariates are the same for all individuals within a population. This makes detecting weather effects more difficult because the signal may be washed out if the temporal (e.g., across months) and spatial (e.g., across a site) aggregation scheme does not match the underlying, and unknown, relationships between weather and demographic rates. We therefore take an alternative, phenomenological approach by modeling size-dependent plant responses to environmental conditions using random year effects. What random year effect models lack in mechanism, they gain in general applicability to a range of ecological problems. The generality stems from the fact that random year effect models essentially use a single index to account for all climate covariates, which also has the added benefit of avoiding over-fit models.

We are interested in whether environmental responses depend on individual size, so we compare two types of vital rate models. The first GLMM assumes that environmental responses are independent of size, meaning that random year effects are applied only to model intercepts. The second GLMM assumes that environmental responses vary depending on individual size, meaning that random year effects are applied to both model intercepts and to the effect of individual size (e.g., a random intercepts, random slopes model). At their simplest, our vital rate statistical models take the form:

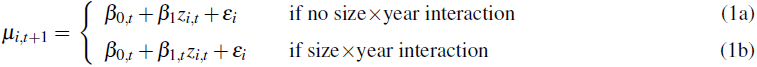

where *μ*_*i,t*+1_ could represent predicted individual size^1^ or the logit of survival probability for individual *i* at time *t* + 1, *z*_*i,t*_ is the size of individual *i* at time *t*, β_0,*t*_ is the intercept for year *t*, β_1_ is the effect of individual size, and ***e*** are i.i.d. errors. The only difference between the two models is that the effect of size, β_1_, includes a subscript *t* in eqn. (1b) that allows the effect to vary across years when we wish to explicitly model a size ×; year interaction. Without the *t* subscript (eqn. 1a), the effect of size is the same every year. Models with and without size ×; year interactions can then be compared using likelihood ratio tests.

### An empirical example: perennial plant populations

To demonstrate our approach, we use long-term data for fifteen perennial plant species from five semi-arid grasslands (Chu & Adler, 2015). Each site includes a set of 1-m^2^ permanent quadrats within which all individual plants were identified and mapped annually using a pantograph (Hill 1920). The resulting mapped polygons represent basal cover for grasses and canopy cover for shrubs. Data come from the Sonoran desert in Arizona (Anderson *et al.*, 2012), sagebrush steppe in Idaho (Zachmann *et al.*, 2010), southern mixed prairie in Kansas (Adler *et al.*, 2007), northern mixed prairie in Montana (Anderson *et al.*, 2011), and Chihuahuan desert in New Mexico (Chu & Adler, 2015). Demographic data on plant growth, survival, and recruitment was extracted using the computer algorithm described by Lauenroth & Adler (2008), Chu *et al.* (2014), and Chu & Adler (2015). The time span of the data range from 13 to 38 year-to-year transitions. The data include observations of likely seedlings and individuals of age 1, which are all given the same arbitrarly small size. We removed these observations before conducting the statistical analysis below to avoid biasing our results toward these individuals of unknown, but very small, size. We fit a separate model describing the probability of a seedling transitioning to a known size for our integral projection model (SI Section SI.1)

We modeled the survival probability and growth of an individual genet as a function of genet size, permanent spatial location (group of quadrats), the crowding experienced by the focal genet from conspecific neighbors, and temporal variation among years. We fit models with and without a size-by-year interaction for each vital rate. The survival probability (*S*) of genet *i* in quadrat group *g* from time *t* to *t* + 1 is estimated as:

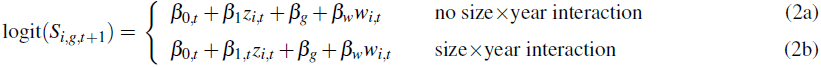

where β_0,*t*_ is the intercept for year *t*, β_1,*t*_ is the effect of log plant size (*z*_*i,t*_) in year *t*, β_*g*_ is the intercept offset for quadrat group *g*, and β_*w*_ is the effect of intraspecific crowding (*w*_*i,t*_). Similarly, we model the log size (*z*) of genet *i* in quadrat group *g* at time *t* + 1 as:

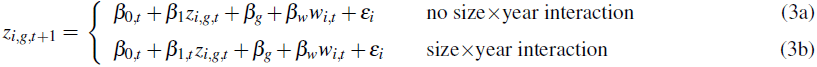

where the ***β***s are as described for the survival model in equation 2. We modeled the intraspecific crowding effect (*w*) based on the size of neighboring conspecifics and their distance from each focal genet (SI Section SI.2). We fit the survival and growth models for each of our 15 species using the lme4∷lmer() function (Bates *et al.*, 2015) in the statistical computing environment R (R Core Team, 2016). We compared models with and without size-by-year interactions for each vital rate and species using likelihood ratio tests based on the expected likelihood ratio distribution under the null hypothesis of no size-by-year interaction. We performed the likelihood ratio tests in this way because standard likelihood ratio tests based on a χ^2^ distribution are not appropriate for comparing random effects structures (SI Section SI.3).

The effect of a size-by-year interaction coefficient on a focal vital rate depends on the size of the individual. Simply comparing the interaction coefficients through time for different sizes does not give us much information on the correlation of responses through time or the variance of those responses. Therefore, to compare large and small individual’s responses to the environment, we calculated growth and survival *anomalies*, which measure how much the size-dependent expected growth rate or survival probability in a given year deviates from the across-year average of the same quantity.

To calculate anomalies for growth, for each year (i.e., each year-specific set of regression coefficients) we calculated the predicted log-size (*z*_*i,t*+1_) of a small individual (10^th^ percentile for the species) and a large individual (90^th^ percentile), assuming an average level of competition 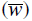 For each individual, we then calculated the difference between the predicted log-size (*z*_*t*+1_) and the original log-size (*z*_*t*_). We then repeated these calculations for the same individuals in an average year (average coefficients). The differences between each year-specific projected size change for year *t*, and the average year projected size change, is defined as the growth anomaly *ℊ*_t_for that individual. Similarly, to calculate the annual survival anomaly (*ℐ*_*t*_), we compared size-specific predicted survival probabilities in each year to the predicted survival in an average year, for large and small individuals.

We used these anomalies to evaluate the correlation of large and small plant responses using the R function cor(x, y, method = “pearson”) and the cor.test() function to calculate *P* values. We also calculated the responsiveness of small and large plants to environmental fluctuations as the standard deviation of the yearly anomalies,σ (*ℊ*_*t*_) and σ (*y*_*t*_). To test whether small and large plants have different responsiveness, we performed two-way Analysis of Variance (ANOVA) using the empirical values of, σ (*ℊ*_*t*_) and, σ (*ℐ*_*t*_) pooled across species treated as data and plant size as the sole predictor. We ran the ANOVA in R, using the lm() function to fit the model and the Anova() function to extract *F* and *P* values. A significant *P*-value (*P <* 0.05) would indicate that small and large plants have different responsiveness to environmental variability. Together, the correlation tests and the responsiveness tests tell us whether paths **a** (negatively covarying sizes) or **b** (reduced variance of one size) in Fig. 1 are possible.

### Statistical results

We found a significant size-by-year interaction in most species-site combinations (Table 1), indicating that size interacts with individual responses to environmental conditions. Survival models with a size ×;year effect received more support than models without a size ×;year effect for 11 out of 15 species. Growth models with a size ×;year effect received more support than models without a size ×;year effect for all 15 species. These statistical results suggest that size-by-environment interactions are common, especially for growth. Size-by-environment interactions may appear more common for growth than survival in part because real-valued responses (growth) have greater statistical precision than binary responses (survival), making interactions easier to detect.

**Table 1:** Results from comparing models with and without a size ×;year effect using likelihood ratio tests as described in the main text. LR_obs_. *>* 95% LRnull indicates a significant size ×;year interaction for the random effects, indicated by bold font for the value of LR_obs_.. Source code: anomaly analysis.R.

| State | Species | Survival |  | Growth |  |
| --- | --- | --- | --- | --- | --- |
| | | 95% $LR_{null}$ | $LR_{obs.}$ | 95% $LR_{null}$ | $LR_{obs.}$ |
| Idaho | ARTR | 2.76 | **13.51** | 4.31 | **91.39** |
| Idaho | HECO | 4.43 | 2.63 | 4.43 | **68.95** |
| Idaho | POSE | 4.16 | 1.39 | 4.43 | **23.14** |
| Idaho | PSSP | 3.81 | **13.59** | 5.10 | **110.90** |
| Montana | BOGR | 3.86 | **225.51** | 4.48 | **50.07** |
| Montana | HECO | 3.61 | **30.87** | 4.39 | **37.77** |
| Montana | PASM | 3.84 | **61.79** | 4.42 | **156.57** |
| Montana | POSE | 3.89 | **18.46** | 5.17 | **44.02** |
| Arizona | BOER | 4.55 | 1.83 | 3.94 | **5.79** |
| Arizona | BORO | 4.02 | **9.52** | 4.75 | **110.94** |
| New Mexico | BOER | 4.34 | **4.46** | 4.36 | **26.12** |
| New Mexico | SPFL | 4.21 | **9.66** | 4.62 | **16.47** |
| Kansas | ANGE | 4.46 | **9.15** | 4.83 | **21.98** |
| Kansas | BOCU | 4.03 | **7.67** | 4.66 | **31.33** |
| Kansas | BOHI | 3.72 | 1.12 | 4.68 | **21.59** |

The next question is whether demographic variations at different sizes negatively covary, or if particular sizes are less sensitive to environmental variation through time. We addressed these questions using our computed annual anomalies. The year-specific anomalies for large (90^th^ percentile of the empirical size distribution) and small (10^th^ percentile) plants of each species tended to be positively correlated (Figure 2A-B). Cases where small and large plants had opposing responses were relatively rare (SI section SI.4). Only two species had marginally significant negative correlations between small and large plant responses: survival anomalies for large and small *Bouteloua gracilis* (BOGR, Montana) were negatively correlated (Pearson’s *ρ*= *-*0.66, *P* = 0.014) and growth anomalies for large and small *Pascopyrum smithii* (PASM, Montana) were negatively correlated (Pearson’s *ρ*= *-*0.60, *P* = 0.029). Variance of year-specific anomalies was similar between small and large plants for growth (*F*_1,28_ = 0.241, *P* = 0.627) but different for survival (*F*_1,28_ = 12.11, *P* = 0.002) when pooling all species (Fig. 3). Thus, we found little evidence for negative covariances between size-based anomalies (path **a** in Fig. 1), but we did find evidence for large plants having reduced variance of survival relative to small plants (path **b** in Fig. 1). This finding is consistent with previous work based on matrix population models (Morris *et al.*, 2008).

**Figure 2:**
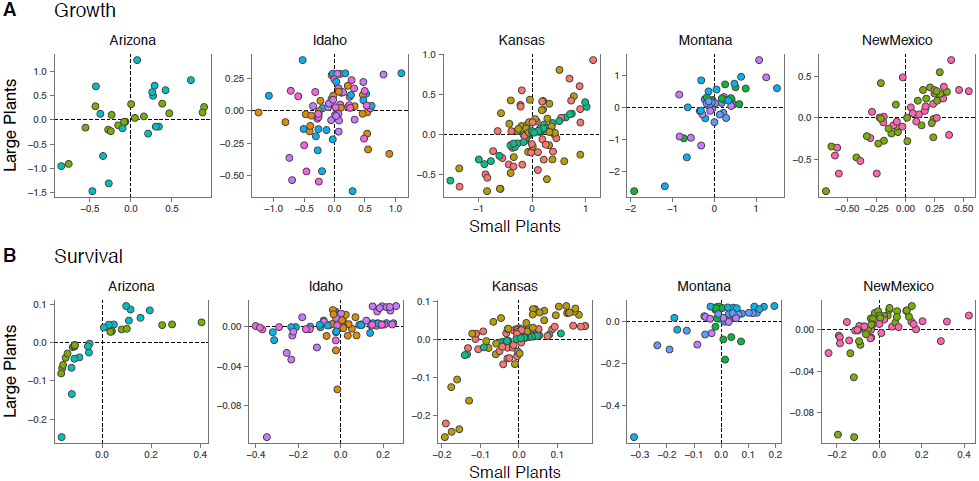
Scatterplots of annual growth (*ℊ*_*t*_) and survival (*ℐ*_*t*_) anomalies by state (panels) and species (colors). (**A**) Relationship between small and large plant growth anomalies (*ℊ*_*t*_). (**B**) Relationship between small and large plant survival anomalies (*ℐ*_*t*_). Negative or null correlations indicate the potential for demographic buffering via size-related tradeoffs. Each point is an annual anomaly associated with a random year effect for the intercept and the size effect. Note the change in *x*and *y*-axis scalings across panels. Source code: plot sxy statistics.R, anomaly analysis.R.

## Impact of Size-by-Environment Interactions on Population Dynamics

Statistically significant size-by-environment interactions imply interesting biology: differently sized individuals have distinct responses to environmental variation. But whether size-by-environment interactions impact population dynamics depends on how they are mediated by elasticities and nonlinearities in model functional forms, which reflect a combination of biology and statistical choices. For example, dramatic environmental responses for certain size-vital rate combinations might catch our attention, but they may not impact population growth. Alternatively, seemingly small differences among sizes for a given vital rate (e.g., slightly lower variance of large individual survival) can have large impacts on population dynamics. In the next two sections, we discuss how elasiticities and model functional form combine to determine the influence of individual-level responses on population dynamics.

**Figure 3:**
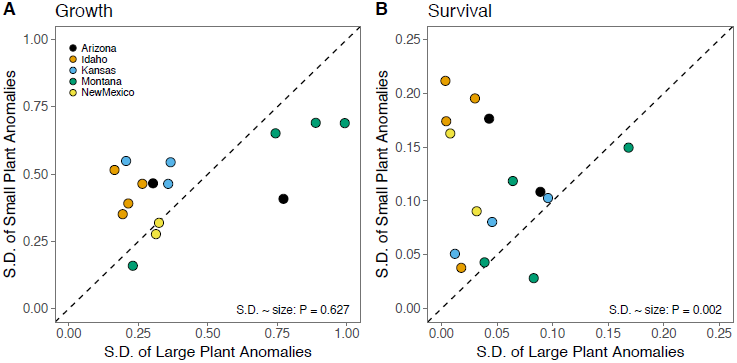
Within-species variation across years in growth (**A**) and survival (**B**) using standard deviations of annual anomalies [σ(*ℊ*_*t*_) and *s (ℐ*_*t*_)] from models with size-dependent environmental responses. Each point is the standard deviation of the yearly anomalies for a given species. Colors represent different sites and the dashed line is 1:1. Source code: plot sxy statistics.R, anomaly analysis.R.

### Elasticities

Elasticities of population models tell us which vital rates and sizes most impact population dynamics. Elasticities quantify the proportional change in population growth rate that results from a proportional change in demographic processes such as survival, growth, and recruitment (de Kroon *et al.*, 1986). In the context of size-by-environment interactions, the relevant elasticities are the elasticity of stochastic population growth rate (λ_*S*_) to between-year variance at different locations in the projection kernel or matrix.

To demonstrate this, we return to our empirical example of 15 perennial plant populations. We used the fitted growth and survival regressions from above, along with a regression for plant recruitment at the quadrat scale, to build a size-structured integral projection model (IPM) for each species (SI section SI.5). Seedlings for some of our species had distinct vital rates, different from what would be predicted in a strictly size-based model fitted to all plants. We therefore used a two-stage, size-structured IPM for four of the fifteen species (SI section SI.5). We used survival and growth regressions with the size ×;year effect (eqns. 2b and 3b).

Elasticities showed that larger plants are most important for population growth for most species (Fig. 4). However, small plants were important for *Pseudoroegneria spicata* (PSSP, Idaho), *Pascopyrum smithii* (PASM, Montana), and *Poa secunda* (POSE, Montana) (Fig. 4). For all species, the survival-growth kernel creates a ridge of high elasticity, indicating survival and growth are most important for population growth in these perennial species, with little contribution from recruitment. Our statistical analysis showed that large plants have reduced variance in survival relative to small plants. Combined with the elasticity results, this could indicate demographic buffering through selection for reduced sensitivity of large plant survival. However, whether reduced variance would be selected for or against depends on the norm of reaction between the environment and the focal vital rate.

### Model nonlinearities

As mentioned above (**Size-by-environment interactions and population dynamics**), stochastic matrix model and IPM theory suggest that demographic variability in matrix entries or kernel values will always decrease population growth rate, and that selection therefore will always favor decreased variance (i.e., demographic buffering). But demographic variance can, in theory, increase fitness and population growth rates because of nonlinear averaging (Rees *et al.*, 2004; Koons *et al.*, 2009).

**Figure 4:**
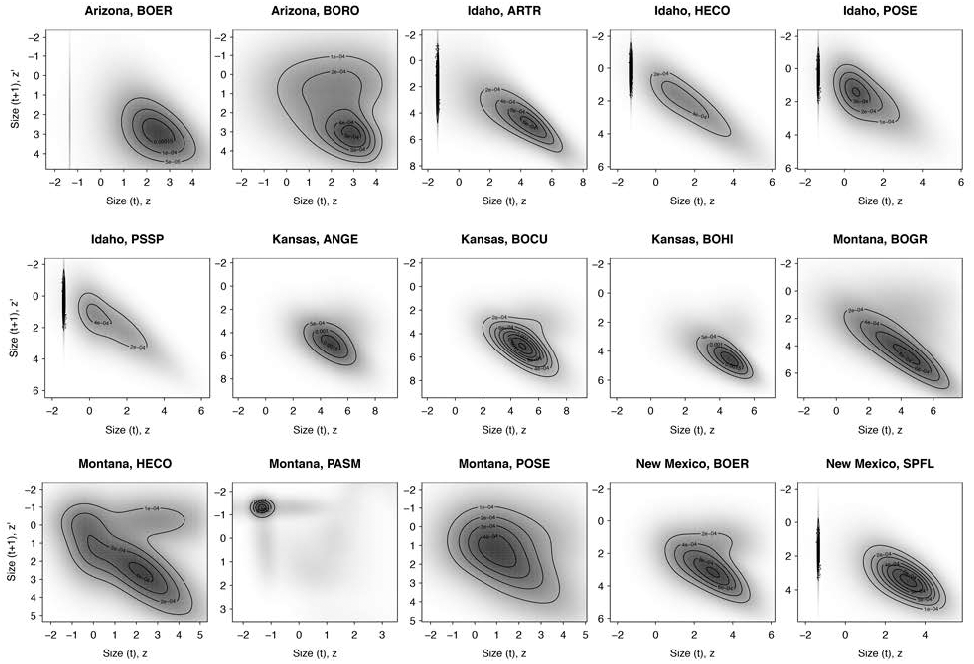
Elasticities for each species. The greyscale surface is on the squareroot scale to increase visibility. Contours are on the arithmetic scale. Two-stage IPMs can be identified by the vertical ridge at small values of size (*z*) on the x-axis. Source code: plot sensitivities.R, sensitivity analysis.R.

A nonlinear average is the average of a nonlinear function of some variable over that variable’s distribution, and is typically different from the value of the function at the *mean* of the distribution. For example, if metabolic rate is a nonlinear function of temperature, metabolic rate at the average temperature is likely to differ from the metabolic rate averaged over many instances of observed temperature. Whether the nonlinear average is greater or less than the value at the mean of the variable quantity depends on the shape of the function (Jensen, 1906), which is often called the *norm of reaction* to the variable. Much previous work has focused on norms of reaction between population growth rate (λ) and environmental variables (*x*) such as temperature (Drake, 2005). If the function is concave-up, the mean and variance of λwill increase as the variance of *x* increases. Concave-up norms of reaction do not automatically imply that environmental fluctuations are beneficial – it depends on the relative impacts on the arithmetic mean and variance of λover time, though increasing log-concave up relationships tend to increase geomtric means (Cohen, 1980; Drake, 2005). If the function is linear or concave-down, the arithmetic mean of λdecreases, and the variance of λincreases, as the variance of *x* increases. This is always bad for population growth because decreasing the arithmetic mean and increasing the variance of population growth through time both act to reduce the geometric mean ofλ, which represents long-run population growth rate (λ_*S*_).

The possible impacts (positive and negative) of environmental fluctuations on population growth are consequences of Jensen’s inequality (Jensen, 1906), and lead to two evolutionary hypotheses. First, linear and concave-down norms of reaction between population growth rate and the some environmental variable should select for demographic buffering, where the most important vital rates vary less than other vital rates in fluctuating environments (Morris *et al.*, 2008). Second, concave-up norms of reaction can select for demographic lability, where the most important vital rates track fluctuating environments (Koons *et al.*, 2009).

As we have shown statistically, size-by-environment interactions allow different size individuals to have more or less variability over time (Fig. 3). Analagous to the situation with vital rates, whether such reduced sensitivity to environmental fluctuations has a positive, negative, or null impact on population growth rate depends on the reaction norms between individual-level vital rates and the environment. However, our statistical models do not include explicit relationships between vital rates and the environment because we used random year effects to approximate the effect of a temporally fluctuating environment. The relevant reaction norm is therefore the implicit one between the year effect and each vital rate (SI Section SI.6). For growth, the reaction norm between the year effect and growth is concave-up because we fit the growth model based on the log of plant area. This is because the exponentiated growth function is concave-up with respect to the year effects (Fig. SI-1A). If including size-by-year interactions in addition to year effects on the intercept alters the distribution of expectations of year-specific growth, then size-by-year interactions might benefit population growth.

For survival, the reaction norm between the year effect and survival is concave-up before the inflection point in the logistic function – i.e., for predicted survival 0.5 or lower – and concave-down after the inflection point (Fig. SI-1B). Therefore, including size-by-year interactions in addition to year effects on the intercept can be beneficial for population growth under the following conditions: (i) the size-by-year effects alter the distribution of survival expectations and (ii) small sizes (with low survival) are more variable. In contrast, size-by-year interactions can hurt population growth under the following conditions: (i) the size-by-year effects alter the distribution of survival expectations and (ii) large (with high survival) sizes are more variable. Note that, for both growth and survival, size-by-year interactions will have no impact on population growth if the interactions do not alter the distribution of year-specific expectations.

Given our model functional forms and our elasticity surfaces (Fig. 4), we might expect increased variance of large plant growth to benefit population growth (because of the concave-up reaction norm between growth and the random year effects) and increased variance of large plant survival to negatively impact population growth (concave-down reaction norm). Similarly, we might expect high survival variance of small plants, which we observe, to have a positive impact population through demographic lability (concave-up reaction norm).

To test these predictions, we used the IPMs for each species in simulation experiments where we manipulated the variance of either large or small individuals in response to environmental fluctuations. The starting point of the analysis are the fitted growth and survival regressions without size ×;year interactions. We then varied the size-dependence of interannual variability, while keeping the total amount of variability constant (SI Section SI.7). We then calculated the stochastic low-density growth rate for each species from the IPM for three experiments: (1) size-independent year effects, (2) size-dependent year effects with large plants more variable than small plants, and (3) size-dependent year effects with small plants more variable than large plants.

In many cases, making small or large plants more variable had little impact on population growth (Fig. 5). But where the scenarios did make an impact, the impacts match our expecations. Making large plant growth more variable increased population growth rates by as much as 20% (Fig. 5), reflecting the benefit conferred to population growth by the concave-up reaction norm between the environment and growth. Perturbing survival variance had much less impact than perturbing growth (Fig. 5). Thus, for our focal species, size-by-environment interactions ca impact population growth through large plant variance in individual growth. Our empirical models, however, showed that large plant growth is not more variable than small plant growth. We did find a difference in variance for survival between large and small plants, but, as our buffering experiments show, altering survival variance among sizes has little impact on population growth.

**Figure 5:**
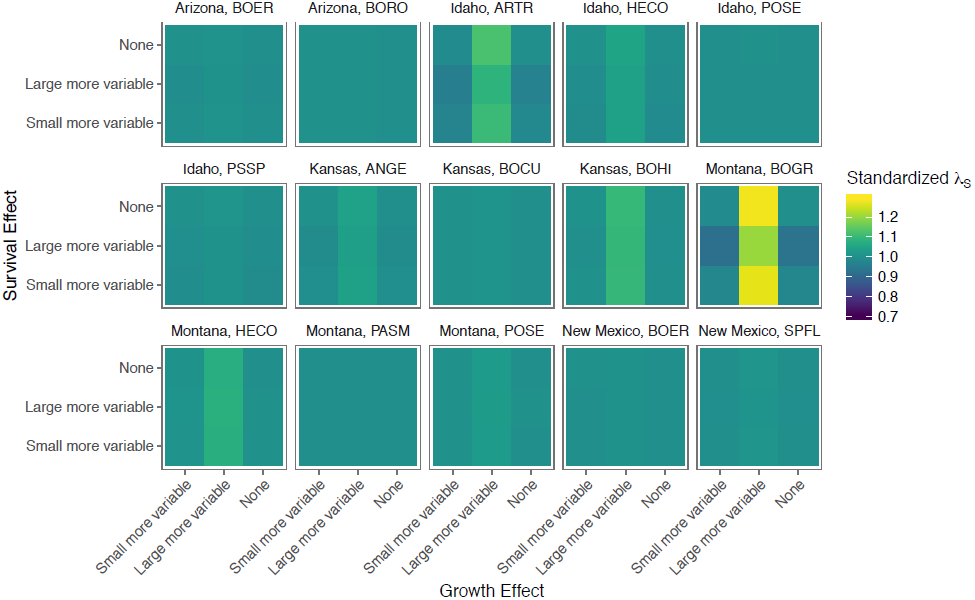
Results from buffering experiments. Standardized stochastic population growth rate (λ_*S*_) is the estimated stochastic lowdensity growth rate from each simulation divided by the stochastic low-density growth rate from the “none” simulations (i.e., no size ×;year interactions). Source code: plot buffering experiments.R, ipm no overlap sxy onestage lambda.R, ipm no overlap sxy twostage lambda.R.

### Population simulations

The elasticity and buffering analyses make it clear that the observed size-by-environment interactions will not impact population dynamics for our focal species. That is, if we have two population models, one with vital rate models with Size ×;year effects and one with random effects only on the intercepts, the stochastic steady-state plant cover from the two models will probably be similar. Although the high survival variance for small plants and low survival variance for large plants that we estimate (Fig. 3) are in line with evolutionary expectations stemming from Jensen’s inequality, our elasticity analysis shows that small plants are not very important for population growth (Fig. 4) and perturbing the size-specific variance of survival has little impact on population growth (Fig. 5). Therefore, the benefit of high survival variance for small plants will not impact population-level dynamics. As a sanity check, however, we can perform simulations to validate these arguments.

We simulated the IPM to estimate the stochastic equilibrium and temporal variance of cover for each species with and without the size ×;year effect. We also use the IPM to simulate transient dynamics after a perturbation to small size classes, allowing us to test whether including the size ×;year interaction impacts the return time to equilibrium cover. In all of our simulations, we set the random effects due to quadrat group to zero, meaning that we are simulating dynamics on a hypothetical average plot. We refer to this as a “hypothetical average plot” because the relationship between quadrat group effect and demographic response is also nonlinear, so setting the group effect to zero is not a way to estimate average cover across groups. We used kernel selection (Metcalf *et al.*, 2015) to incorporate temporal environmental variation, meaning that at each time step we randomly selected model coefficients corresponding to one observation year for the survival, growth and recruitment models.

For equilibrium runs, we initialized species with very low cover and ran the model for 500 time steps (years) to allow the community to reach its steady-state pattern of fluctuations in response to environmental variability. We calculated the average steady-state cover 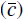 for each species by simulating an additional 2000 time steps and averaging cover over this period. We also stored the entire 2000 year simulation for each species to examine its distribution. Equilibirum runs were conducted with and without the size ×;year interaction.

For transient runs, we initialized each species with very low cover composed of only small plants and ran the model for 100 time steps, long enough for each species to reach average steady-state cover (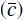 as calculated from equilibrium runs). We did this 50 times for each species, and for each run calculated the number of iterations it took to reach average steady-state cover. We then averaged those number of iterations across the 50 simulations to estimate mean return time to equilibrium.

The results are a direct consequence of the fact that the sizes whose variance change most when we model size-by-environment interactions have low elasticity. The larger variation of small plant survival in response to environmental conditions (Fig. 3) had no impact on simulated equilibrium cover or the temporal variance of cover (Fig. 6A). Likewise, transient dynamics, defined as return time to equilibrium after a perturbation to small size classes, were not affected by the size ×;year effect, except for *Artemisia tripartita* (ARTR, Idaho) (Fig. 6B). Return time to equilibrium took longer with the size ×;year effect (mean over 50 simulations = 32.4 years) than without it (mean over 50 simulations = 20.6 years), but these means are associated with high variability (error bars in Fig. 6B).

**Figure 6:**
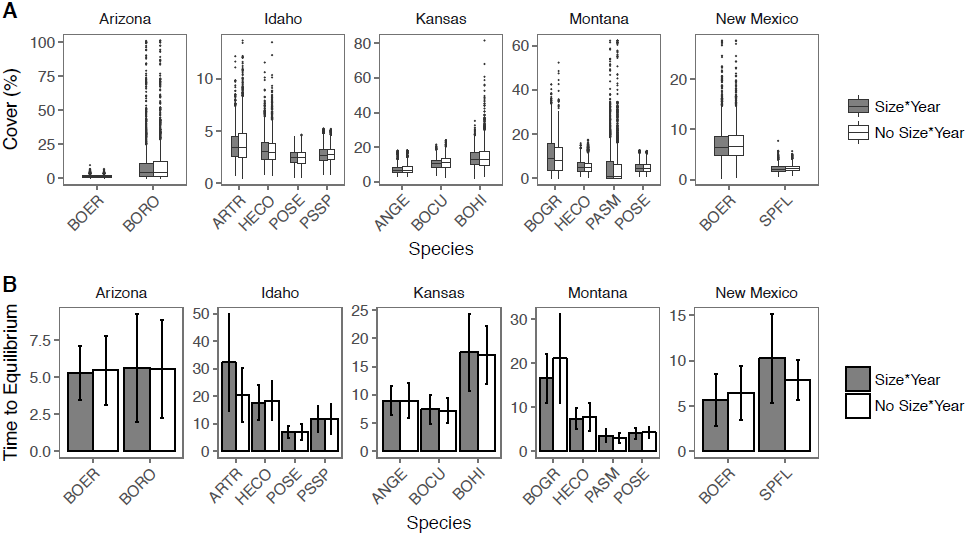
Distribution of plant cover (**A**) and transient dynamics (**B**) from IPM simulations with and without a size year effect. Boxplots in **A** show the median cover (solid horizontal line), the 25^th^ and 75^th^ percentiles (the box), the highest and lowest values no further than 1.5 ×;the interquartile range (whiskers), and extreme values beyond the whiskers (points). In **B**, “Time to Equilibrium” is the mean (over 50 simulations) return time from very low cover to equilibrium cover. Error bars represent the minimum and maximum time to equilibrium from the 50 simulations. Source code: analyze ipm results.R.

## Discussion

A central focus of population ecology has been quantifying how species respond to environmental fluctuations as a consequence of vital rate-environment relationships (Lewontin & Cohen, 1969; Lawson *et al.*, 2015). Many populations are size-structured and empirical evidence is mounting that different size individuals often have different responses to the environment (*e.g.*, Morris *et al.*, 2008; Ozgul *et al.*, 2010; Zhang *et al.*, 2011; Lindmark *et al.*, 2018). Just as different vital rates may have negative covariances (Morris & Doak, 2004), environmental responses by individuals of different sizes may negatively covary, or some sizes may be less sensitive to envrionmental fluctuations than others (Fig. 1). Whether such size-by-environment interactions have positive or negative population-level consequences depends on the norms of reaction between environment variables and the demographic responses (Fig. 1). How these reaction norms align with size-dependent elasticities determines the population-level impacts of size-by-environment interactions.

### Size-by-environment interactions are common

Our case study of 15 plant populations from five semiarid grasslands showed that statistically significant size-by-year interactions are common (Table 1). Assuming that the variability modeled by random year effects in our regression models represent the species’ response to fluctuating environmental conditions, these statistical results suggest that size-by-environment interactions are a potentially common phenomenon. Stage- and age-structured matrix population models include (st)age-by-environment interactions when elements of the projection matrix are allowed to vary through time (*e.g.,* Morris *et al.*, 2008), but whether each element should be allowed to vary through time is rarely tested statistically. Our statistical approach can be easily applied to age-structured populations because age is a continuous trait.

Because they may be common, it is important to conduct statistical tests for size-by-year interactions whether or not they ultimately impact population dynamics. Statistically significant size-by-year interactions help reduce unexplained variance in models of vital rates, while also representing more faithfully the variability in data (Clark, 2003). Assuming all individuals share a common response to interannual variation (i.e., no size ×;year interaction) can misrepresent the variability of the data by producing estimates of random year effects that do not actually apply to any particular size. For example, averaging over distinct responses of small and large individuals could result in parameter estimates that fall somewhere in between the two real responses. This could leave substantial unexplained variance that can cause problems when models also include explicit time-varying covariates because too much residual variation can be attributed to the covariate effects^2^. To avoid these potential pitfalls, we suggest demographic modelers test for size-by-year interactions using our mixed-effects framework.

The strength of the mixed-effects modeling approach is that it does not require estimating separate parameters for individual size, stage, or age classes. Likewise, our approach is general because we assume random year effects represent the response of species and sizes to interannual variation in the environment. Thus, our approach can be applied to any dataset with multiple years of data, and the basic demographic models are all generalized linear models. Our use of random year effects is also the main weakness of our approach because we cannot attribute environmental responses to particular environmental drivers and thus cannot predict responses to environmental change. But given the difficulty and number of years of data required for identifying correctly the environmental drivers of population dynamics and selecting among competing models (Teller *et al.*, 2016), we think that our phenomenological approach will often be a good compromise between ignoring size-dependent responses and the ideal situation where size-specific responses are linked to identified time-varying environmental covariates. However, fitting random year effects requires making statistical choices that can bias model-based inference, as we discuss below (*The importance of model functional form*).

### Population level impacts of size-by-year interactions

We found that small and large plants tended to respond similarly to temporal environmental fluctuations (Fig. 2). This indicated that negative covariance among sizes’ responses to the environment did not occur, and therefore could not lead to demographic buffering. The other route to population-level effects, size-dependent magnitude of sensitivity to environmental fluctuations, was possible because large plant survival (where variability is harmful) was less variable than small plant survival (where variability is helpful, Figs. 3 and SI-1).

Ultimately, our empirical results do not suggest demographic buffering or demographic lability in our study populations, despite statistically significant size ×;year effects. Demographic buffering or lability requires that the size being acted upon be important for population dynamics. Because small plants have low elasticity for population growth rate, natural selection for lability should be weak for small plants, as implied by our statistical results and model functional forms. On the contrary, large plants have high elasticity, making them candidates for strong natural selection. However, we found no evidence of high growth variance for large plants, which would be beneficial for population growth according to our analysis. Thus, several ingredients for evolution of demographic buffering or lability via size-byenvironment interations were present in our case studies, but they did not all align to impact population dynamics. In part, our “negative” results are due to the fact that including size-by-environment interactions did not alter the average expectation across years for small or large plants, for both growth and survival (Figs. SI-3 and SI-4). Whether our results are generalizable depends on if our results are system specific.

If our results are system specific, then they might not be generalizable beyond similar species (semi-arid perennial bunchgrasses). Semi-arid plants may not be particularly sensitive to environmental fluctuations due to bet-hedging, in which case adding size-dependence to models of temporal variability would not have a large impact (Figs. SI-3 and SI-4). Moreover, the difference of environmental responses between small and large perennial bunchgrasses might be small despite being statistically significant. Species with larger average differences in size between small and large individuals might display larger differences in environmental responses, just as the magnitude of trait differences at the species-level corresponds to differences in environmental responses among species (Polis, 1984; Angert *et al.*, 2009). Species with larger ranges of size than our focal species might exhibit stronger size-by-year interactions. For example, stage-structured models of forests treat seedlings and adult trees as essentially different species because their vital rates and the environment they experience (extreme shade for seedlings, high light for adult trees) are so different (e.g., Freckleton *et al.*, 2003). As a result, the response of seedlings and adult trees to environmental fluctuations are probably also distinct, and their differences are likely more extreme than the differences between small and large bunchgrass individuals. Size-by-environment interactions may still prove to be important in other systems and investigating size-by-environment interactions could reveal what kinds of species exploit environmental fluctuations and what kinds of species merely cope with environmental fluctuations.

If, on the contrary, our results are generalizable to other systems because population dynamics are not generally affected by size-dependent responses to the environment, it would be in spite of the prevalence of size-by-environment interactions. Several of the other potential mechanisms by which populations may buffer against environmental variability have been examined, including reduction of the variance of sensitive vital rates and correlations among vital rates (Jongejans *et al.*, 2010; Lawson *et al.*, 2015; Compagnoni *et al.*, 2016; McDonald *et al.*, 2017). Across studies and mechanisms, empirical evidence suggests that some populations seem buffered against variability, while others do not. It is also true that scientists (including ourselves) tend to study species where individuals exist in abundance (Crone *et al.*, 2013). It then stands to reason that population biologists also tend to study species under environmental conditions where their populations persist, perhaps even despite measurable size-dependent responses and/or a lack of buffering. These artifacts of study design are not necessarily uninformative; instead it may suggest that, under most circumstances, normal environmental fluctuations should not be expected to strongly affect populations where individuals exist in abundance. More research is required to show whether size-dependent responses have more significant population-level impacts at range centers versus range edges, or under changing conditions.

Here we have examined plant populations, but our core ideas and the analytical approach apply to animal populations, too. Size-by-environment interactions may be especially influential in animal populations because of complex ontogeny in animals. For example, certain fish and reptiles span four or more orders of magnitude in body weight over the course of development (reviewed in Werner & Gilliam, 1984), making young (small) and adult (large) individuals within a population as dissimilar as individuals from different species (*e.g.,* Polis, 1984). Likewise, Bieber & Ruf (2005) showed that the importance of different life-stages for population growth of wild boar (*Sus scrofa*) shift depending on environmental conditions. These lines of evidence suggest size-by-environment interactions may be common in animal populations.

### The importance of model functional form

Are our results actually indicative of species’ life histories? Or, are our results and conclusions pre-determined by our choice of model functional forms? We log-transformed individual size in our growth models in order to obtain a simple model: linear, with variance that was not strongly size-dependent. We also compared the log transform with square root, third root, and other transformations before settling on log transformation. Log transformation is appealing because it reduces heteroscedasticity, and growth is typically a multiplicative biological process. When growth is determinate (i.e., expected growth is positive below some size, and negative above that size), a linear growth model on log scale automatically results in absolute growth rate (on arithmetic scale) being maximized at intermediate sizes, and relative growth rate decreasing with size, patterns that are commonly observed in nature (Ellner *et al.*, 2016, Ch. 2). But another unavoidable consequence is that the growth model is concave-up as a function of random intercept and slope coefficients, which we assume represent environmental fluctuations, in the linear predictor of the growth regression (SI Section SI.6). Any subsequent model-based inquiry within our mixed-models framework as to whether a species is better off with higher or lower sensitivity to environmental fluctuations at a given size is constrained, by Jensen’s inequality, to one possible answer: higher variability increases mean growth rate.

A similar “one possible answer” situation is present in our survival model, a GLMM with the conventional sigmoidal (logit) link function for fitting to binomial data. A logistic regression for survival as a linear function of size *z*, is concave-up for survival *<* 0.5, concave-down for survival *>* 0.5, and approximately linear near the inflection point. Therefore, variability in random effects coefficients for survival will be beneficial for small plants, which tend to have lower survival rates in the concave-up portion of the link function, and harmful for large plants which tend to have higher survival rates in the concave-down portion of the model. If survival in an average year is near 1, then a good year cannot help much, but a bad year can hurt.

To disentangle true size-dependent (or size-independent) responses to environmental variability from effects of model functional form, modellers need to pay special attention to the curvature of demographic responses to environmental variability. Standard practices such as log-transforming individual size or logistic regression for survival should be checked against alternatives, and against more flexible modeling approaches such as generalized additive models (e.g., Wood, 2000; Ramsay *et al.*, 2009), which have the potential to capture both positive and negative curvature where they exist (Rees *et al.*, 2014). Another possibility is to choose different distributions for the random effects (e.g, assume random year effects vary around the mean following a lognormal rather than normal distribution). However, flexible regression modeling functions are not a cure-all. The built-in criteria and algorithms for choosing model complexity (like conventional model selection criteria such as AIC) try to identify optimal models for predicting the response, not optimal models for predicting the curvature of the response (e.g., Fan & Gijbels, 1996), and they often achieve parsimony by biasing estimates towards a simple conventional model such as logistic or linear regression. As a result, default smoothing methods can seem to give you an answer about curvature when the real situation is “not enough data.” For example, if a model of *y* as a linear function of *x* receives as much support as a model of *y* as a linear function of 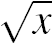, then the real answer is “not enough data.” Exploratory data analyses, such as plotting average survival or average growth rates for binned subsets of the population, may be the most informative approach.

## Conclusions

Size-by-environment interactions might be an important way for species to cope with and/or exploit environmental variation. Or, they might not matter much in terms of population growth rate, despite being biologically interesting. Indeed, we found that size-by-environment interactions are common in perennial plant species, but they have little impact on population dynamics. Given the vast literature on size-based differences among intraspecific individuals, it is likely that size-by-environment interactions are common in many other species. Unfortunately, it is difficult to guess whether size-by-environment interactions, if present, will tend to impact population dynamics in other species. This is because of the many contingencies involved: reaction norms must be nonlinear, size-by-environment interactions must impact sizes and vital rates with high elasticity, and size-by-environment interactions must alter the variance of individual-level vital rates. Only future research in other systems can tell if our results are generalizable.

Our case studies did bring into stark relief how statistical choices can impact model-based inference. In particular, the link functions used in our vital rate models inherently introduce nonlinearity in vital rate responses to environmental variability, when environmental variability is represented as random year effects. The functional forms of vital rate models represent a mixture of biology and convenience. On the convenience side, modeling growth on log-transformed scale makes for a well-behaved model, as does assuming that random year effects have a normal distribution centered on zero, rather than some other mean effect of the environment that might be nonzero. On the biological side, individual growth *is* a multiplicative process and patterns of increased variance with size are common in nature. Making progress will require new developments that allow demographic modellers to select vital rate models based on their ability to predict a response as well as the curvature of that response (Ye & Hooker, 2018). We hope that this paper serves as a catalyst for these developments.

## Acknowledgments

We thank Robin Snyder for helpful discussions and comments on the manuscript, and we thank David Koons for comments on an earlier version of the manuscript. This research was supported by U.S. National Science Foundation grants DEB-1353078 (USU) and DEB-1353039 (Cornell).

## Authorship Statement

All authors conceived the ideas and designed methodology. ATT and BJT conducted the analyses. ATT and BJT wrote the first draft of the manuscript, and all authors contributed substantially to revisions.

## Supporting Information

### Section SI.1 Are seedlings different?

We excluded likely seedlings from our analysis of size-(in)dependent environmental responses, but we need to include seedlings in our IPMs to accurately reproduce population dynamics. Seedlings are only partially identified in the original quadrat maps from which our demographic data is derived. So for the IPM analyses presented in the main text, likely seedlings were identified by size and age for fitting vital rate models as follows. An individual recorded as age=1 in the original data is apparently being identified as a seedling. However, such individuals in a quadrat that was not observed the previous year are actually of unknown age. If they are small, such individuals might be seedlings or they might be older individuals that shrank to a seedling-like size. So for all analyses of seedlings only, or of older individuals only, we removed from the data set all such “doubtful” individuals: small (size ≤0.25 cm^2^) and recorded as age=1 in a quadrat that was not observed the previous year. With doubtful individuals removed, any remaining individual who is small (size≤0.25 cm^2^) and recorded as age=1 are classified as seedlings, and all others are classified as older. This assumes that any individual larger than 0.25 cm^2^ not a seedling, regardless of their recorded age.

To test whether seedlings are demographically different, we fit survival models that included a factor variable flagging likely seedlings. A significantly nonzero coefficient for this variable implies that seedlings behave differently, all else being equal. Specifically, we fitted the model

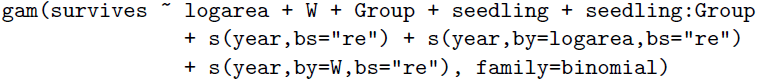

where *W* is within-species competition and Group is the quadrat group. The model includes random year effects on the intercept, the slope in size, and the slope in *W*.

The results from these model fits are shown in Table SI-1. All the Idaho species have seedlings that are different, requiring the two-stage IPM described in the main text. Likewise, *B. eriopoda* in Arizona and *S. flexuosus* in New Mexico also require a two-stage IPM.

**Table SI-1:**
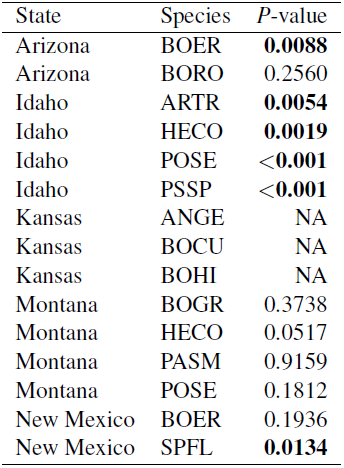
Results from fitting GAMMs for survival with a seedling effect. Shown here are just the *P*-values for the seedling effect. Kansas has no genets that get flagged as seedlings.

| State | Species | $P$-value |
| --- | --- | --- |
| Arizona | BOER | **0.0088** |
| Arizona | BORO | 0.2560 |
| Idaho | ARTR | **0.0054** |
| Idaho | HECO | **0.0019** |
| Idaho | POSE | **<0.001** |
| Idaho | PSSP | **<0.001** |
| Kansas | ANGE | NA |
| Kansas | BOCU | NA |
| Kansas | BOHI | NA |
| Montana | BOGR | 0.3738 |
| Montana | HECO | 0.0517 |
| Montana | PASM | 0.9159 |
| Montana | POSE | 0.1812 |
| New Mexico | BOER | 0.1936 |
| New Mexico | SPFL | **0.0134** |

### Section SI.2 Modeling local crowding

We modeled the crowding experienced by a focal genet in each year *t* as a function of the distance to and size of neighbor genets. In previous work, we assumed that the decay of crowding with neighbor distance followed a Gaussian function (Chu & Adler, 2015), but here we use a data-driven approach (Teller *et al.*, 2016). We model the crowding experienced by genet *i* of species *j* from it conspecific neighbors as the sum of neighbor areas across a set of concentric annuli, *k*, centered at the plant,

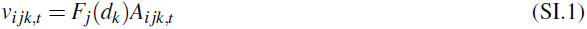

where *F*_*j*_ is the competition kernel (described below) for effects of intraspecific crowding on species *j, d*_*k*_ is the average of the inner and outer radii of annulus *k*, and *A*_*ijk,t*_ is the total area of genets of species *j* in annulus *k* around the focal genet in year *t*. The total crowding on the focal genet exerted by its own species is

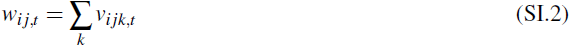

The *w*’s are then included as covariates in the survival and growth regressions. Note that we drop the subscript *j* in our equations for vital rates for clarity because *inter*specific competition is not considered in this paper.

We assume that competition kernels *F*_*j*_(*d*) are non-negative and decreasing, so that distant plants have less effect than close plants. Otherwise, we let the data dictate the shape of the kernel by fitting a spline model using the methods of Teller *et al.* (2016). The shape of *F*_*j*_ is determined by a set of spline basis coefficients 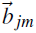 and a smoothing parameter η that controls the complexity of the fitted kernel. Demographic models such as our survival model,

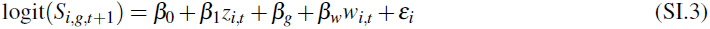

then have ɛ, **β**, 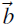 and ηas parameters to be fitted. We implemented this in the statistical computing environment, R (R Core Team, 2016), by making the spline coefficients and η the arguments of an objective function that calculates each *v*_*ijk,t*_ value using the input spline coefficients, calls the model-fitting functions lmer and/or glmer (Bates *et al.*, 2015) to fit the other parameters in the survival and growth regressions, and returns an approximate AIC value and model degrees of freedom (*df*) for survival and growth combined. We used the 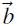 values at the smoothest (smallest *df*) local minimum of AIC as a function of *df*, as in Teller *et al.* (2016). This approach assumes that one measure of crowding affects survival and growth.

Once we had estimated the competitions kernels, we used them to calculate the *w*_*ijk,t*_ values for each individual in each year, and fitted the full survival and growth regressions, which include the interspecific interaction coefficients, ω(see main text). All genets in a quadrat were included in calculating the *w*’s, but plants located within 5 cm of quadrat edges were not used in fitting the regressions.

### Section SI.3 Likelihood ratio tests for size-by-year interactions

We compared models with and without size-by-year interactions for each vital rate and species using likelihood ratio tests based on the expected likelihood ratio distribution under the null hypothesis of no sizeby-year interaction. We performed the likelihood ratio tests in this way because standard likelihood ratio tests based on a χ^2^distribution are not appropriate for comparing random effects structures This is because the reduced model is on the boundary of the set of parameter values for the full model – it is at σ= 0 for one of the variance components (Scheipl *et al.*, 2008). Specifically, we simulated responses from the fitted models without size-by-year interactions (Eqs. 2a, 3a), used those values as data to refit models with (Eqs. 2b, 3b) and without (Eqs. 2a, 3a) a size-by-year interaction, and then calculated the likelihood ratio between those two fits. By repeating this procedure 500 times, we generated the likelihood ratio distribution under the null hypothesis of no size-by-year interaction. We then compared the observed likelihood ratio to the 95^th^ quantile of the null distribution to perform a one-sided significance test. If the observed likelihood ratio exceeds the 95^th^ quantile of the null distribution, then the size-by-year interaction is statistically significant.

### Section SI.4 Descriptive statistics from anomaly analysis

#### Section SI.4.1 Tally of anomalies with equal signs: comparing large and small plants

In the output below, offs refers to the number of annual anomalies that have different signs (plus or minus) for small and large plants, ons refers to the number of annual anomalies that have the same sign (plus or minus) for small and large plants, and off perc and off perc are those tallies represented as percentage of the number of anomalies.

~~~
state species vital offs ons off_perc on_perc
1 Arizona BOER grow 2 14 12.500000 87.50000
2 Arizona BOER surv 0 17 0.000000 100.00000
3 Arizona BORO grow 5 12 29.411765 70.58824
4 Arizona BORO surv 0 17 0.000000 100.00000
5 Idaho ARTR grow 8 13 38.095238 61.90476
6 Idaho ARTR surv 11 10 52.380952 47.61905
7 Idaho HECO grow 6 15 28.571429 71.42857
8 Idaho HECO surv 5 16 23.809524 76.19048
9 Idaho POSE grow 9 12 42.857143 57.14286
10 Idaho POSE surv 5 16 23.809524 76.19048
11 Idaho PSSP grow 6 15 28.571429 71.42857
12 Idaho PSSP surv 3 18 14.285714 85.71429
13 Kansas ANGE grow 14 24 36.842105 63.15789
14 Kansas ANGE surv 11 27 28.947368 71.05263
15 Kansas BOCU grow 18 19 48.648649 51.35135
16 Kansas BOCU surv 6 31 16.216216 83.78378
17 Kansas BOHI grow 1 33 2.941176 97.05882
18 Kansas BOHI surv 5 30 14.285714 85.71429
19 Montana BOGR grow 4 9 30.769231 69.23077
20 Montana BOGR surv 11 2 84.615385 15.38462
21 Montana HECO grow 2 11 15.384615 84.61538
22 Montana HECO surv 4 9 30.769231 69.23077
23 Montana PASM grow 10 3 76.923077 23.07692
24 Montana PASM surv 1 12 7.692308 92.30769
25 Montana POSE grow 2 11 15.384615 84.61538
26 Montana POSE surv 4 9 30.769231 69.23077
27 NewMexico BOER grow 7 23 23.333333 76.66667
28 NewMexico BOER surv 8 22 26.666667 73.33333
29 NewMexico SPFL grow 7 22 24.137931 75.86207
30 NewMexico SPFL surv 10 20 33.333333 66.66667
~~~

Grouped by state and vital rate,

~~~
state vital offs ons off_perc on_perc
1 Arizona grow 7 26 21.21212 78.78788
2 Arizona surv 0 34 0.00000 100.00000
3 Idaho grow 29 55 34.52381 65.47619
4 Idaho surv 24 60 28.57143 71.42857
5 Kansas grow 33 76 30.27523 69.72477
6 Kansas surv 22 88 20.00000 80.00000
7 Montana grow 18 34 34.61538 65.38462
8 Montana surv 20 32 38.46154 61.53846
9 NewMexico grow 14 45 23.72881 76.27119
10 NewMexico surv 18 42 30.00000 70.00000
~~~

Grouped by vital rate,

~~~
vital offs ons off_perc on_perc
1grow 101 236 29.97033 70.02967
2 surv 84 256 24.70588 75.29412
~~~

#### Section SI.4.2 Tally of anomalies with equal signs: comparing growth and survival

In the output below, offs refers to the number of annual anomalies that have different signs (plus or minus) for growth and survival, ons refers to the number of annual anomalies that have the same sign (plus or minus) for growth and survival, and off perc and off perc are those tallies represented as percentage of the number of anomalies.

state species plant_size offs ons off_perc on_perc

~~~
1 Arizona BOER Large Plants 1 15 6.25000 93.75000
2 Arizona BOER Small Plants 3 13 18.75000 81.25000
3 Arizona BORO Large Plants 7 10 41.17647 58.82353
4 Arizona BORO Small Plants 8 9 47.05882 52.94118
5 Idaho ARTR Large Plants 8 13 38.09524 61.90476
6 Idaho ARTR Small Plants 3 18 14.28571 85.71429
7 Idaho HECO Large Plants 11 10 52.38095 47.61905
8 Idaho HECO Small Plants 8 13 38.09524 61.90476
9 Idaho POSE Large Plants 10 11 47.61905 52.38095
10 Idaho POSE Small Plants 10 11 47.61905 52.38095
11 Idaho PSSP Large Plants 8 13 38.09524 61.90476
12 Idaho PSSP Small Plants 9 12 42.85714 57.14286
13 Kansas ANGE Large Plants 24 14 63.15789 36.84211
14 Kansas ANGE Small Plants 17 21 44.73684 55.26316
15 Kansas BOCU Large Plants 11 26 29.72973 70.27027
16 Kansas BOCU Small Plants 13 24 35.13514 64.86486
17 Kansas BOHI Large Plants 17 17 50.00000 50.00000
18 Kansas BOHI Small Plants 17 17 50.00000 50.00000
19 Montana BOGR Large Plants 5 8 38.46154 61.53846
20 Montana BOGR Small Plants 4 9 30.76923 69.23077
21 Montana HECO Large Plants 2 11 15.38462 84.61538
22 Montana HECO Small Plants 4 9 30.76923 69.23077
23 Montana PASM Large Plants 3 10 23.07692 76.92308
24 Montana PASM Small Plants 8 5 61.53846 38.46154
25 Montana POSE Large Plants 3 10 23.07692 76.92308
26 Montana POSE Small Plants 5 8 38.46154 61.53846
27 NewMexico BOER Large Plants 13 17 43.33333 56.66667
28 NewMexico BOER Small Plants 10 20 33.33333 66.66667
29 NewMexico SPFL Large Plants 12 17 41.37931 58.62069
30 NewMexico SPFL Small Plants 9 20 31.03448 68.96552
~~~

Grouped by state and vital rate

~~~
state plant_size offs ons off_perc on_perc
1 Arizona Large Plants 8 25 24.24242 75.75758
2 Arizona Small Plants 11 22 33.33333 66.66667
3 Idaho Large Plants 37 47 44.04762 55.95238
4 Idaho Small Plants 30 54 35.71429 64.28571
5 Kansas Large Plants 52 57 47.70642 52.29358
6 Kansas Small Plants 47 62 43.11927 56.88073
7 Montana Large Plants 13 39 25.00000 75.00000
8 Montana Small Plants 21 31 40.38462 59.61538
9 NewMexico Large Plants 25 34 42.37288 57.62712
10 NewMexico Small Plants 19 40 32.20339 67.79661
~~~

Grouped by vital rate

~~~
plant_size offs ons off_perc on_perc
1 Large Plants 135 202 40.05935 59.94065
2 Small Plants 128 209 37.98220 62.01780
~~~

### Section SI.5 Integral projection model details

#### Section SI.5.1 Modeling recruitment

Unlike the survival and growth models, our recruitment model does not include an effect of individual size because a new recruit has no size at time *t –* 1 by definition. Therefore, the recruitment model does not include a size ×;year interaction. Following previous work (Chu & Adler, 2015), we model recruitment at the level of the quadrat becuase the data do not attribute new genets to specific parents. We assume the number of recruits (*y*) in quadrat *q* in a given year follows a negative binomial distribution, whose mean is a function of the cover of the species, quadrat group, temporal variation among years, and intraspecific interactions as a function of species’ cover within a quadrat. The recruitment model is:

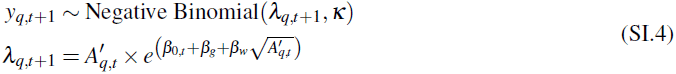

where λ is the mean number of recruits occuring across space, *к* is the dispersion parameter, the β s are as in the survival and growth regressions, and *A*^*′*^ is the effective cover of the focal species. We define “effective cover” as a mixture of observed cover in the focal quadrat, *q*, and the mean cover, 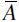 across the group, *g*, in which the focal quadrat is located:

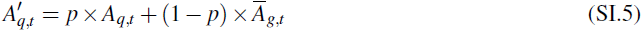

where *p* is a mixing fraction between 0 and 1, estimated by the model.

We fit the recruiment model in JAGS (Just Another Gibbs Sampler) (Plummer, 2003), using the rjags package (Plummer, 2014) to connect JAGS and R. Posterior distributions of all unknown parameters were sampled from three parallel MCMC (Monte Carlo Markov Chain) chains that ran for 20,000 iterations, thinned by 50, after an initial burn-in of 10,000 iterations. We assessed convergence of MCMC chains visually and by ensuring the multivariate scale reduction factor was less than 1.1 (Gelman & Rubin, 1992). The multivariate scale reduction factor was calculated using the coda∷gelman.diag() function (Plummer *et al.*, 2006).

#### Section SI.5.2 Survival and growth models for seedlings

We excluded seedlings from the data for fitting the statistical models above because they would have biased our tests of whether individual size interacts with interannual environmental variability. However, when we move to modeling species’ populations with the IPM, we will need information on the seedling stage: what is the probability that a seedling survives? What is the probability that a seedling transitions to a non-seedling size class, and what is the distribution of possible sizes? Our model for seedling survival is similar to that of non-seedlings, except we exclude the effect of genet size because all seedlings have the same size. The model for seedling survival (lowercase *s*) is:

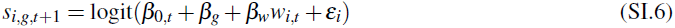

where the terms are as in equation 2 in the main text.

Our model for seedling growth combines two probabilities: (1) the probability that a seedling transitions to a non-seedling size class and (2) the probability of being a given non-seedling size. We use a mixture of two Guassians, which combine a left peak of seedlings and a right peak of non-seedlings. The seedling peak has mean = log(0.25) and standard deviation = 0.2 by fiat. The distribution parameters of the non-seedling peak are the fraction in the left peak (*f*), and the mean (*μ*) and standard deviation (σ) of the size of age 2 seedlings that became large enough to measure. The mixture distribution takes the form: *f* ×;N log(0.25), 0.2^2^ +(1 – *f*) ×;N (*μ*,σ^2^). All parameters were estimated by maximum likelihood using the stats4∷mle(…, method = ‘‘Nelder-Mead’’) function in R (R Core Team, 2016).

#### Section SI.5.3 Integral projection models for perennial plants

We used two single species IPM structures, depending on whether seedlings were identified as behaving differently than non-seedlings for a given species. Seedlings may have lower probability of survival than non-seedlings, on average, meaning we would need to separate the life stages in those cases. To test whether seedlings are different from non-seedlings, we fit survival models where genets were flagged as seedling or non-seedling based on their recorded age and size. The seedling factor was included in the survival model for each species. If the seedling factor was significant (*P <* 0.05), we concluded that seedlings were different and used a two-life stage model for those species. We first describe the one-stage IPM, which is similar to our previous work (Chu & Adler, 2014; Tredennick *et al.*, 2017), and then describe the two-stage IPM.

In the one-stage IPM, the size distribution of a species is represented by a density function *n(u,t*) which gives the density of genets of size *u* at time *t*. The size distribution function at time *t* + 1 is given by

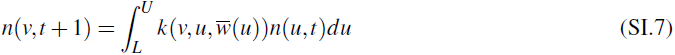

where 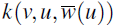 is the population kernel that describes all possible transitions from size *u* to *v* and 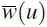 is a scalar representing the average intraspecific crowding experienced by a genet of size *u*. The integral is evaluated over all possible sizes between predefined lower (*L*) and upper (*U*) size limits that extend beyond the range of observed genet sizes. The kernel 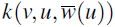 is constructed from our fitted survival (*S*), growth (*G*), and recruitment (*R*) regression models.

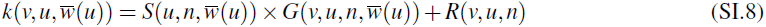

The recruitment regression predicts the number of recruits per unit area, and we assume that the number of recruits increases linearly with genet size, *R(v, u, n*)= exp(*u)R(v, u, n*), to incorporate recruitment in the IPM (Chu & Adler, 2015).

Adler *et al.* (2010) developed a mean field approximation for local crowding when the competition kernels are all Gaussian functions,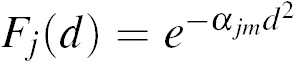. The approximation is explained in the online SI to Adler *et al.* (2010) and in section 5.3 of Ellner *et al.* (2016). Here we explain how that approximation was modified for the IPMs in this paper, which used fitted nonparametric competition kernels (Adler *et al.*, 2018). The mean field approximation is based on the observed spatial distribution patterns of the species (Chu & Adler, 2015). In the observed data conspecific individuals displayed non-random, size-dependent patterns: small genets were randomly distributed, while large genets were segregated from each other. The overdispersion of large conspecific genets is incorporated into the IPM with a ‘no-overlap’ rule, as in (Adler *et al.*, 2010).

We assume that conspecifics cannot overlap (‘no-overlap’ rule). Genet shapes are irregular, but we nonetheless implement the no-overlap rule by assuming that a genet of log area *u*_*i*_ is a circle of radius *r*_*i*_ where 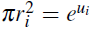. The no-overlap rule is then that the centroids of two conspecific individuals must be separated by at least the sum of their radii.

For any one focal genet, the no-overlap restriction on its neighbors’ locations affects only a negligibly small part of the habitat. The expected cover of individuals in the places where they can occur (relative to one focal plant) is thus assumed to equal their expected locations in the habitat as a whole.

Let *C*_*j*_(*u*) be the total cover of species *j* genets of radius *r* or smaller,

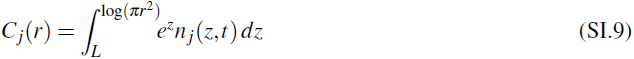

A focal genet of radius *r* cannot have any conspecific neighbors centered at distances less than *r*. It can have a neighbor centered at distance *x > r* if that neighbor’s radius is no more than *x – r*. Adding up the expected cover of all such possible neighbors for a focal genet of radius *r*,

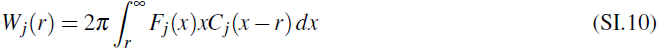

This integral is finite and computable because the kernels *F* fall to 0 at finite *x*.

Our two-stage IPM is exactly the same as the one-stage IPM for non-seedlings whose log(area) *>* 0.25, but includes separate dynamics for seedlings, whose size we set by definition to log(0.25) (see **Survival and growth models: seedlings**). In the two-stage IPM, the size distribution of a species is still represented by a density function *n(u,t*). But the population kernel (*k*) is composed of two parts, one for seedlings and one for non-seedlings:

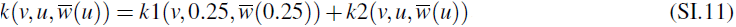

where *k*1 is the seedling kernel defining all possible transitions from the seedling size (0.25) to non-seedling sizes and *k*2 is the non-seedling population kernel as in equation SI.8. The seedling population kernel is defined as 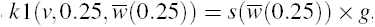, where *s* is the seedling survival regression (eq. SI.6) and *g* is the mixture distribution for seedling growth (see **Survival and growth models: seedlings**). Table SI-1 indicates for which species we use the one-stage IPM and for which we use the two-stage IPM.

### Section SI.6 Year effect reaction norms

Here we provide additional details on the relationship between vital rates, size, and the environment. As mentioned in the main text, the reaction norm of interest for our vital rate statistical models is the implicit one between the vital rate and the random year effects. We need to know whether the variability among year-specific responses results in a concave-up, concave-down, or parabolic reaction norm to guide our inference on the impacts of year effects and size-by-year interaction effects on population growth. We discuss our growth model first, and then discuss the survival model.

#### Section SI.6.1 Year effect reaction norms: growth

Recall the growth model with random year effects on the intercept, aimed at predicting size (*z*) at time *t*,

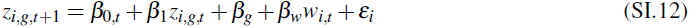

where we can rewrite β_0,*t*_ as β_0_ + *E*_*t*_ such that β_0_ is the mean intercept and *E*_*t*_ is the random effect of year *t*. Then we can rearrange Eq. SI.12 to express growth (*G* = *z*_*t*+1_ *z*_*t*_) of a given individual in real size (area, rather than log area), ignoring quadrat group effects and intraspecific crowding, as

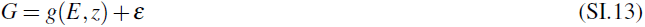

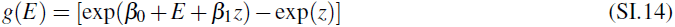

where *z* is log size, β_0_ is the mean intercept, *E* is the random effect (*E* = “environment”), and ε is random error, meaning that *g(E*) is concave-up as a function of *E* (Fig. SI-1A). To make things concrete, the expectation of a log-normal(*μ*, σ) is exp(*μ* + σ^2^*/*2). Or specifically, if you fit a random effect with mean zero on log-transformed scale, this is equivalent to multiplication by a random positive number on arithmetic scale, and that positive number has mean exp(σ^2^*/*2). I.e., modifying Eq. SI.14

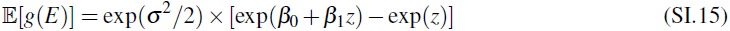

We can introduce random effect specification into this model in several equivalent ways. Here we use the conventional approach of specifying random effects in the same units as the linear predictor, yielding Jensen’s inequality after we re-exponentiate.

The reaction norm between growth and the environment is also concave-up when we include size-byyear interactions, where

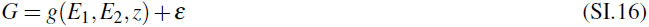

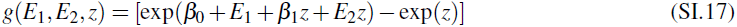

and the second derivatives of *g* with respect to *E*_1_ and *E*_2_ are positive,

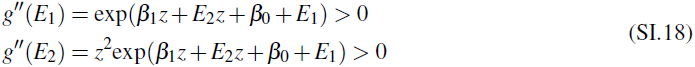

#### Section SI.6.2 Year effect reaction norms: survival

Recall the survival model with a random year effect on the intercept

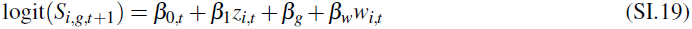

where we can again rewrite β_0,*t*_ as β_0_ + *E*_*t*_ such that β_0_ is the mean intercept and *E*_*t*_ is the random effect for year *t*. Then for a given individual of size *z*, we can rewrite Eq. SI.19 with the antilogit transform applied as

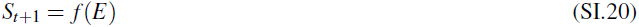

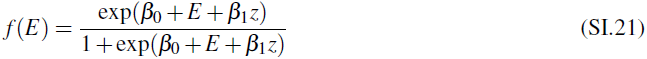

At a given size *z*, the second derivative of *f* with respect to *E* is

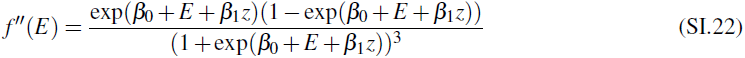

*f* is therefore concave-up (*f*^″^ *>* 0) if and only if exp(β_0_ + *E* + β_1*z*_) *<* 1. This is equivalent to the predicted survival probability *S* being less than 0.5, the value of the response at the inflection point in the logistic function *e*^*u*^*/*(1+ *e*^*u*^), which is in turn equivalent to the linear predictor in the logistic regression, β_0_ + *E* + β_1*z*_, being less than 0. Thus, the reaction norm between year effects and survival is concave-up before the inflection point, and concave-down after the inflection point (Fig. SI-1B). The same is true when we include a size ×;year interaction.

**Figure SI-1:**
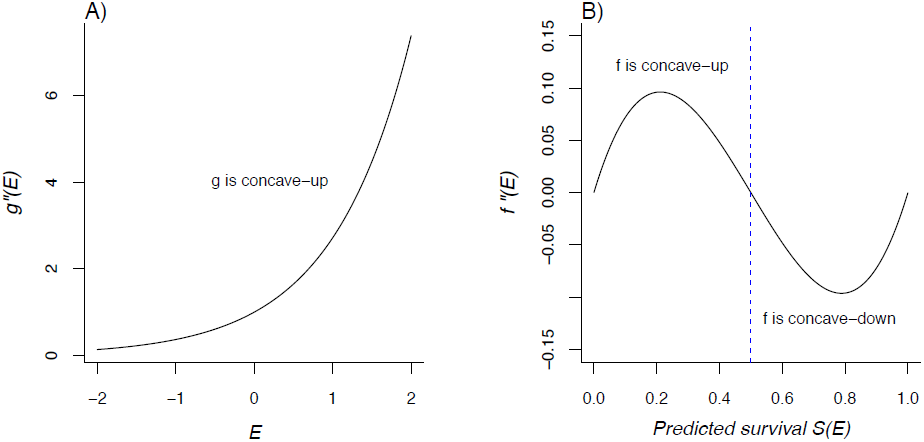
Concavity of the implicit reaction norms between the random year effects (*E*) and the vital rates, growth (A) and survival (B). In B, the dashed blue line shows the location of the inflection point (0.5); concave-up reaction norms occur where *f*^″^(*E*) *>* 1 and concave-down reaction norms occur where *f*^″^(*E*) *<* 1. Source code: reaction norms.R.

### Section SI.7 Vital rate buffering experiments

The starting point is a demographic model with a size-independent year effect,

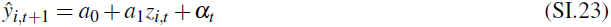

Here ŷ_*i,t*+1_ is the predicted mean response for individual *i* in the subsequent year *t* + 1, *z*_*i,t*_ is that individual’s size in year *t*, α_*t*_ is the year effect for year *t*. In our models, ŷcould be logit survival, the mean of the conditional size distribution, or expected production of new recruits.

We assume that the average of α_*t*_ over years is exactly 0. Depending on the fitting method, this might be true or it might just be approximately true. But it can always be made true, by subtracting the mean from each α_*t*_ and adding it to the intercept *a*_0_.

We want to vary the size-dependence of interannual variability, while keeping the total amount of variability constant. We measure “total amount” from the perspective of individuals, as follows. Let *Z* = {*z* _*j*_, *j* = 1, 2,*, N*} be the set of all sizes of all individuals observed in the data set, regardless of year. The *anomaly a*_*j,t*_ for individual *j* in year *t* is the difference between the value of ŷ_*j,t*+1_ including year-specific random effects, and what ŷ_*j,t*+1_ would be if all year-specific random effects in the model were set to zero. The RMS anomaly *A* is the root-mean-square all *a*_*j,t*_*j* values,

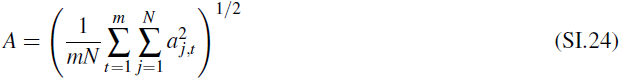

Note that an anomaly is defined for each size ×;year combination: we put every observed size in every observed year, and ask what the anomaly would be. The reason for doing this is that the IPM is simulated by drawing years at random from those in the data set. So for matching the IPM to the true level of between-year variability, our measure of the “true level” should be one in which years are weighted equally, rather than weighted by sample size.

For the baseline model (SI.23), in a given year all sizes have the same anomaly, so the RMS anomaly is

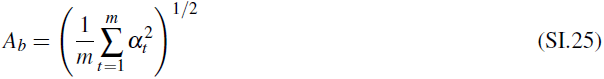

Figure SI-2 shows three size year models that can be tuned to have the same RMS anomaly as the baseline model.

#### Small are more variable

First, we make small plants more variable than large plants. To do this, we choose a size 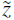 representing a typical plant. We use an exponential tapering of variance so that we don’t have to worry about lines crossing (instead of converging) at sizes representing large and small plants.

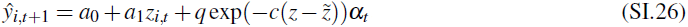

The anomalies are 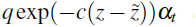 is determined by computing the RMS anomaly with *q* = 1, and then adjusting *q* so that the RMS anomaly becomes *A*_*b*_. The value of *c >* 0 determines how much difference there is between large and small plant variability. A plant of size 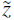 has the same anomalies as in the baseline model.

#### Large are more variable

Second, we make large plants more variable than small plants. This looks the same as above, except that the tapering goes the other way,

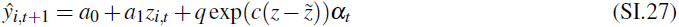

with *c >* 0.

**Figure SI-2:**
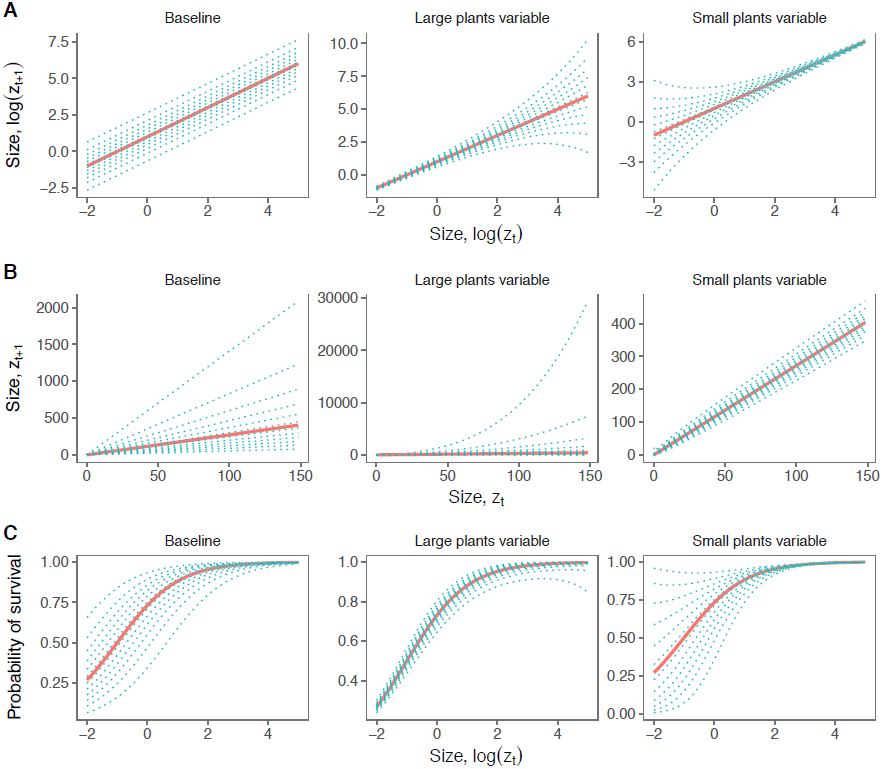
Red lines are the responses with year effect set to 0. Blues lines are responses in different years, such that the root mean square anomaly is the same in all plots. In (A), regressions are shown on the log scale. In (B), the same regressions are shown on the arithmetic scale. In (C), the response is logit-transformed to demonstrate the effect of size-by-environment interactions on survival regressions. Note the difference in scales on the y-axes across panels. Source code: BufferingExperiments.R.

### Section SI.8 Additional figures

**Figure SI-3:**
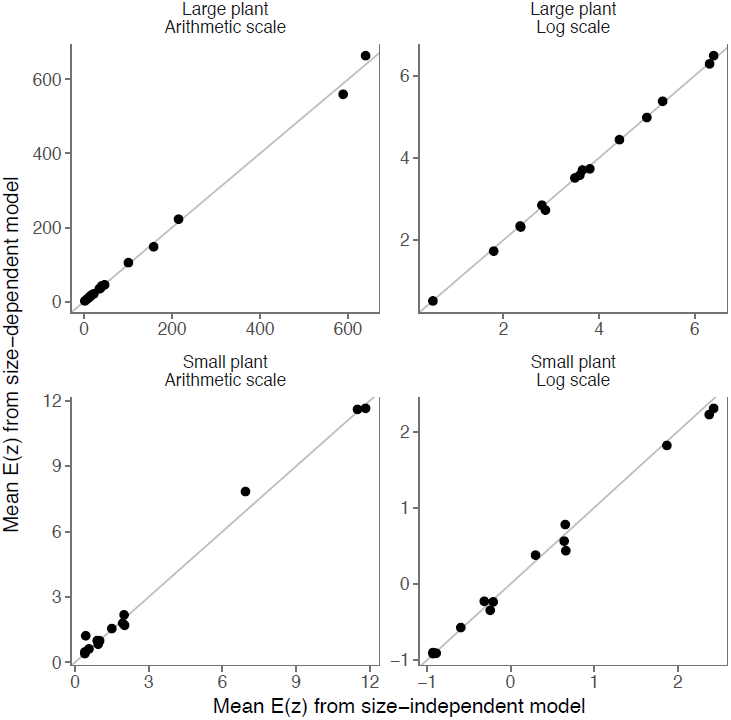
Comparison of growth expectations, averaged over year-specific expectations, from models with and without size ×;year interactions for each species (15 points), in both log space and arithmetic space. The mean expectations are very similar, meaning that adding size ×;year interactions will not induce a Jensen’s effect beyond that already present due to the reaction norm between the year effects and growth.

**Figure SI-4:**
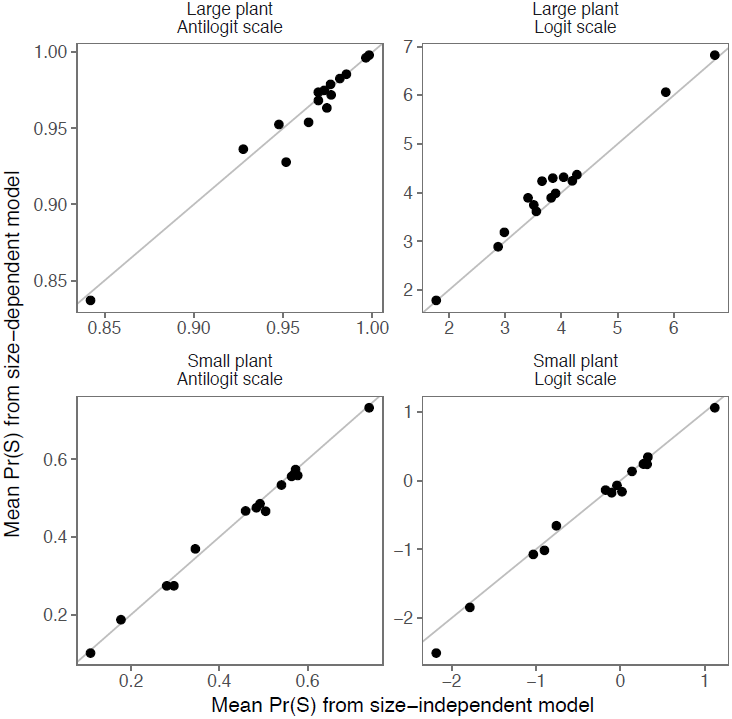
Comparison of survival expectations, averaged over year-specific expectations, from models with and without size ×;year interactions for each species (15 points), in both logit and antilogit space. The mean expectations are very similar, meaning that adding size ×;year interactions will not induce a Jensen’s effect beyond that already present due to the reaction norm between the year effects and survival.

## Footnotes

1 “Size” is often a log-transformed measurement, such as log of a plant’s diameter or ölog of an animal’s body weight.

2 We have seen this in our own work, where including random year effects diminished the importance of an external weather covariate.

## Literature Cited

Adler, P.B. & Drake, J.M. (2008). Environmental variation, stochastic extinction, and competitive coexistence. The American Naturalist, 172, 186–195.

Adler, P.B., Tyburczy, W.R. & Lauenroth, W.K. (2007). Long-term mapped quadrats from Kansas prairie: demo- graphic information for herbaceous plants. Ecology, 88, 2673.

Anderson, J., McClaran, M.P. & Adler, P.B. (2012). Cover and density of semi-desert grassland plants in permanent quadrats mapped from 1915 to 1947. Ecology, 93, 1492–1492.

Anderson, J., Vermeire, L. & Adler, P.B. (2011). Fourteen years of mapped, permanent quadrats in a northern mixed prairie, USA. Ecology, 92, 1703.

Angert, A.L., Huxman, T.E., Chesson, P. & Venable, D.L. (2009). Functional tradeoffs determine species coexistence via the storage effect. Proceedings of the National Academy of Sciences, 106, 11641–11645.

Barton, K.E. & Boege, K. (2017). Future directions in the ontogeny of plant defence: understanding the evolutionary causes and consequences. Ecology Letters, 20, 403–411.

Bates, D., Maechler, M., Bolker, B. & Walker, S. (2015). Fitting linear mixed-effects models using lme4.

Bieber, C. & Ruf, T. (2005). Population dynamics in wild boar Sus scrofa: Ecology, elasticity of growth rate and implications for the management of pulsed resource consumers. Journal of Applied Ecology, 42, 1203–1213.

Chesson, P. (2000). Mechanisms of Maintenance of Species Diversity. Annual Review of Ecology and Systematics, 31, 343–366.

Chu, C. & Adler, P.B. (2015). Large niche differences emerge at the recruitment stage to stabilize grassland coexistence. Ecological Monographs, 85, 373–392.

Chu, C., Havstad, K.M., Kaplan, N., Lauenroth, W.K., McClaran, M.P., Peters, D.P., Vermeire, L.T. & Adler, P.B. (2014). Life form influences survivorship patterns for 109 herbaceous perennials from six semi-arid ecosystems. Journal of Vegetation Science, 25, 947–954.

Clark, J.S. (2003). Uncertainty and variability in demography and population growth: A hierarchical approach. Ecology, 84, 1370–1381.

Cohen, J. (1980). Convexity properties of products of random nonnegative matrices. Proceedings of the National Academy of Sciences, 77, 3749–3752.

Compagnoni, A., Bibian, A.J., Ochocki, B.M., Rogers, H.S., Schultz, E.L., Sneck, M.E., Elderd, B.D., Iler, A.M., Inouye, D.W., Jacquemyn, H. & Miller, T.E. (2016). The effect of demographic correlations on the stochastic population dynamics of perennial plants. Ecological Monographs, 86, 480–494.

Coulson, T., Catchpole, E.A., Albon, S.D., Morgan, B.J.T., Pemberton, J.M., Clutton-Brock, T.H., Crawley, M.J. & Grenfell, B.T. (2001). Age, sex, density, winter weather, and population crashes in Soay sheep. Science, 292, 1528–1531.

Crone, E.E., Ellis, M.M., Morris, W.F., Stanley, A., Bell, T., Bierzychudek, P., Ehrle´n, J., Kaye, T.N., Knight, T.M., Lesica, P., Oostermeijer, G., Quintana-Ascencio, P.F., Ticktin, T., Valverde, T., Williams, J.L., Doak, D.F., Ganesan, R., Mceachern, K., Thorpe, A.S. & Menges, E.S. (2013). Ability of matrix models to explain the past and predict the future of plant populations. Conservation Biology, 27, 968–978.

Dalgleish, H.J., Koons, D.N., Hooten, M.B., Moffet, C.A. & Adler, P.B. (2011). Climate influences the demography of three dominant sagebrush steppe plants. Ecology, 92, 75–85.

De Roos, A.M., Persson, L. & McCauley, E. (2003). The influence of size-dependent life-history traits on the structure and dynamics of populations and communities. Ecology Letters, 6, 473–487.

Drake, J.M. (2005). Population effects of increased climate variation. Proceedings of the Royal Society B: Biological Sciences, 272, 1823–1827.

Ellner, S.P., Childs, D.Z. & Rees, M. (2016). Data-driven Modelling of Structured Populations. Springer International Publishing, Switzerland.

Fan, J. & Gijbels, I. (1996). Local polynomial modelling and its applications. Chapman and Hall, London.

Freckleton, R.P., Silva Matos D.M., Bovi, M.L. & Watkinson, A.R. (2003). Predicting the impacts of harvesting using structured population models: The importance of density-dependence and timing of harvest for a tropical palm tree. Journal of Applied Ecology, 40, 846–858.

Gamelon, M., Gaillard, J.M., Gimenez, O., Coulson, T., Tuljapurkar, S. & Baubet, E. (2016). Linking demographic responses and life history tactics from longitudinal data in mammals. Oikos, 125, 395–404.

Gillespie, J. (1977). Natural selection for variance in offspring numbers: a new evolutionary principle. The American Naturalist, 111, 1011–1014.

Huülsmann, S., Rinke, K. & Mooij, W.M. (2011). Size-selective predation and predator-induced life-history shifts alter the outcome of competition between planktonic grazers. Functional Ecology, 25, 199–208.

Hunter, C.M., Caswell, H., Runge, M.C., Regehr, E.V., Amstrup, S.C. & Stirling, I. (2010). Climate change threatens polar bear populations: A stochastic demographic analysis. Ecology, 91, 2883–2897.

Jensen, J.L. (1906). Sur les functions convexes et les ine´gualite´s entre les valeurs moyennes. Acta Mathematica, 30, 175–193.

Jongejans, E., de Kroon, H., Tuljapurkar, S. & Shea, K. (2010). Plant populations track rather than buffer climate fluctuations. Ecology Letters, 13, 736–743.

Knops, J.M.H., Koenig, W.D. & Carmen, W.J. (2007). Negative correlation does not imply a tradeoff between growth and reproduction in California oaks. Proceedings of the National Academy of Sciences, 104, 16982–16985.

Koons, D.N., Pavard, S., Baudisch, A. & Metcalf, C.J.E. (2009). Is life-history buffering or lability adaptive in stochastic environments? Oikos, 118, 972–980.

de Kroon, H., Plaisier, A., van Groenendael, J. & Caswell, H. (1986). Elasticity: The Relative Contribution of Demographic Parameters to Population Growth Rate. Ecology, 67, 1427–1431.

Lande, R., Engen, S. & Saether, B.E. (2003). Stochastic Population Dynamics in Ecology and Conservation. Oxford University Press, New York, New York.

Lauenroth, W.K. & Adler, P.B. (2008). Demography of perennial grassland plants: Survival, life expectancy and life span. Journal of Ecology, 96, 1023–1032.

Lawson, C.R., Vindenes, Y., Bailey, L. & van de Pol, M. (2015). Environmental variation and population responses to global change. Ecology Letters, 18, 724–736.

Lewontin, R.C. & Cohen, D. (1969). On population growth in a randomly varying environment. Proceedings of the National Academy of Sciences, 62, 1056–1060.

Lindmark, M., Huss, M., Ohlberger, J. & Ga˚rdmark, A. (2018). Temperature-dependent body size effects determine population responses to climate warming. Ecology Letters, 21, 181–189.

McDonald, J.L., Franco, M., Townley, S., Ezard, T.H., Jelbert, K. & Hodgson, D.J. (2017). Divergent demographic strategies of plants in variable environments. Nature Ecology and Evolution, 1.

Metcalf, C.J.E., Ellner, S.P., Childs, D.Z., Salguero-Go´mez, R., Merow, C., McMahon, S.M., Jongejans, E. & Rees, M. (2015). Statistical modelling of annual variation for inference on stochastic population dynamics using Integral Projection Models. Methods in Ecology and Evolution, 6, 1007–1017.

Morris, W. & Doak, D. (2004). Buffering of Life Histories against Environmental Stochasticity: Accounting for a Spurious Correlation between the Variabilities of Vital Rates and Their Contributions to Fitness. The American Naturalist, 163, 579–590.

Morris, W.F., Pfister, C.A., Tuljapurkar, S., Haridas, C.V., Boggs, C.L., Boyce, M.S., Bruna, E.M., Church, D.R.,

Coulson, T., Doak, D.F., Forsyth, S., Gaillard, J.M., Horvitz, C.C., Kalisz, S., Kendall, B.E., Knight, T.M., Lee, C.T. & Menges, E.S. (2008). Longevity can buffer plant and animal populations against changing climatic variability. Ecology, 89, 19–25.

Ozgul, A., Childs, D.Z., Oli, M.K., Armitage, K.B., Blumstein, D.T., Olson, L.E., Tuljapurkar, S. & Coulson, T. (2010). Coupled dynamics of body mass and population growth in response to environmental change. Nature, 466, 482–485.

Pfister, C.A. (1998). Patterns of variance in stage-structured populations: Evolutionary predictions and ecological implications. Proceedings of the National Academy of Sciences, 95, 213–218.

van de Pol, M., Bailey, L.D., McLean, N., Rijsdijk, L., Lawson, C.R. & Brouwer, L. (2016). Identifying the best climatic predictors in ecology and evolution. Methods in Ecology and Evolution, 7, 1246–1257.

Polis, G.A. (1984). Age Structure Component of Niche Width and Intraspecific Resource Partitioning: Can Age Groups Function as Ecological Species? The American Naturalist, 123, 541–564.

R Core Team (2016). R: A Language and Environment for Statistical Computing. Vienna, Austria.

Ramsay, J., Hooker, G. & Graves, S. (2009). Functional Data Analysis with R and MATLAB. Springer-Verlag, New York.

Rees, M., Childs, D.Z. & Ellner, S.P. (2014). Building integral projection models: A user’s guide. Journal of Animal Ecology, 83, 528–545.

Rees, M., Childs, D.Z., Rose, K.E., Grubb, P.J., Ellner, S.P., Rees, M., Rose, K.E., Grubb, P.J. & Ellner, S.P. (2004). Evolution of size-dependent flowering in a variable environment: construction and analysis of a stochastic integral projection model. Proceedings of the Royal Society B: Biological Sciences, 271, 425–434.

Sprules, W.G. (1972). Effects of Size-Selective Predation and Food Competition on High Altitude Zooplankton Com- munities. Ecology, 53, 375–386.

Stacey, P.B. & Taper, M. (1992). Environmental variation and the persistence of small populations. Ecological Applications, 2, 18–29.

Teller, B.J., Adler, P.B., Edwards, C.B., Hooker, G., Snyder, R.E. & Ellner, S.P. (2016). Linking demography with drivers: climate and competition. Methods in Ecology and Evolution, 7, 171–183.

Werner, E.E. & Gilliam, J.F. (1984). The Ontogenetic Niche and Species Interactions in Size-Structured Populations. Annual Review of Ecology and Systematics, 15, 393–425.

Wood, S.N. (2000). Modelling and smoothing parameter estimation with multiple quadratic penalties. Journal of the Royal Statistical Society: Series B (Statistical Methodology), 62, 413–428.

Ye, Z. & Hooker, G. (2018). Local Quadratic Estimation of the Curvature in a Functional Single Index Model. ArXiv e-prints.

Zachmann, L., Moffet, C. & Adler, P. (2010). Mapped quadrats in sagebrush steppe: long-term data for analyzing demographic rates and plantplant interactions. Ecology, 91, 3427.

Zhang, R., Jongejans, E. & Shea, K. (2011). Warming increases the spread of an invasive thistle. PLoS ONE, 6.

## Additional References

Adler, P.B., Ellner, S.P. & Levine, J.M. (2010). Coexistence of perennial plants: An embarrassment of niches. Ecology Letters, 13, 1019–1029.

Adler, P.B., Kleinhesselink, A., Hooker, G., Taylor, J.B., Teller, B.J. & Ellner, S.P. (2018). Weak interspecific interactions in a sagebrush steppe? Conflicting evidence from observations and experiments. Ecology, in press.

Chu, C. & Adler, P.B. (2014). When should plant population models include age structure? Journal of Ecology, 102, 531–543.

Chu, C. & Adler, P.B. (2015). Large niche differences emerge at the recruitment stage to stabilize grassland coexistence. Ecological Monographs, 85, 373–392.

Gelman, A. & Rubin, D.B. (1992). Inference from Iterative Simulation Using Multiple Sequences. Statistical Science, 7, 457–472.

Plummer, M. (2003). JAGS: A Program for Analysis of Bayesian Graphical Models Using Gibbs Sampling. In: Proceedings of the 3rd International Workshop on Distributed Statistical Computing (DSC 2003). March. pp. 20–22.

Plummer, M. (2014). rjags: Bayesian graphical models using MCMC.

Plummer, M., Best, N., Cowles, K. & Vines, K. (2006). CODA: Convergence Diagnosis and Output Analysis for MCMC. R News, 6, 7–11.

R Core Team (2016). R: A Language and Environment for Statistical Computing. Vienna, Austria.

Scheipl, F., Greven, S. & Küchenhoff, H. (2008). Size and power of tests for a zero random effect variance or polynomial regression in additive and linear mixed models. Computational Statistics and Data Analysis, 52, 3283–3299.

Tredennick, A.T., Hooten, M.B. & Adler, P.B. (2017). Do we need demographic data to forecast plant population dynamics? Methods in Ecology and Evolution, 8, 541–551.

